# An artificially selected cytokinin–pathway transcription factor balances soybean yield and pathogen resistance

**DOI:** 10.1101/2025.07.04.663232

**Authors:** Qun Ma, Xueting Hu, Qun Yang, Yiwen Gao, Yucheng Liu, Lei Li, Zhixi Tian, Bo Ren

## Abstract

Soybean contributes to sustainable food and feed production, but some modern varieties favor yield at the cost of disease resistance. Using a bottom-up approach, we demonstrate that the RR2b transcription factor activates cytokinin signaling for plant growth and reproductive fitness but represses disease-resistance and root-nodulation pathways. *RR2b* was artificially selected during domestication and improvement, and its expression levels correlate with ATT repeat polymorphisms in its promoter. Most wild soybeans and landraces have either more or less ATT repeats: more lead to weak *RR2b* expression with lower yield and enhanced resistance to soybean bacterial blight, and fewer cause strong *RR2b* expression, higher yield but reduced disease resistance. Elite cultivars instead balance yield and defense via moderate *RR2b* expression and several ATT repeats. This confers contrasting RR2b functionality across haplotypes in response to contrasting selection pressures and offers insights into decoupling trade-offs between yield and disease resistance to enhance crop productivity.

## Introduction

Crop domestication is considered one of the most significant milestones in human history, marking the transition of human society from a hunting– gathering lifestyle to an agricultural–production system. By around 4,000 years ago, humans had successfully domesticated all major crops that are essential to our diets today, including the legume cultivated soybean *Glycine max*, from its wild progenitor *G. soja*^1^. Today, global soybean production reaches approximately 400 million tons per year, accounting for about half of the world’s oilseed production and contributing over a quarter of the world’s feed protein^2^. Soybean fixes atmospheric nitrogen into plant–available nitrate through symbiotic nitrogen fixation, with the potential to fix 15–20 million tons of nitrogen in the soil each year^3^. This dramatically reduces the economic and environmental costs of crop cultivation and enables sustainable agriculture. The soybean industry suffers annual losses exceeding $US25 billion due to pests and diseases. During soybean domestication, a series of favorable traits, such as yield, seed size, seed hardness, non-shattering, and oil content, were artificially selected to meet human needs. However, this also led to a decline in the genetic diversity of soybean populations (i.e. a genetic bottleneck), making it difficult to breed high–yield and disease-resistant varieties^4^.

Identification of adaptive genes controlling domestication–related traits and characterizing their genetic and molecular mechanisms are integral to understanding the dynamic process of crop domestication and often serve as a launch pad for delivery of improved elite cultivars. There are two principal approaches to identify adaptive genes. A top–down approach (genetic mapping of domestication traits) starts with a phenotype of interest and then seeks to identify causal genomic regions by genetic analysis such as genome–wide association study (GWAS) and linkage–disequilibrium mapping. The other is a bottom–up approach (genome scans for footprints of selection), which begins with identification of adaptation signals in a set of genes or a region in the genome using population–genetic methods, and then moving from gene to phenotype by molecular methods^5^. To date, most genes isolated by the top– down approach are so–called ‘low-hanging fruit’, which are single, large–effect loci controlling major domestication traits, such as *fw2.2* for fruit size in tomato^6^, *Sh4* for non–shattering in rice^7^, *tb1* for plant and inflorescence architecture in maize^8^, *Q* for threshability in wheat^9^, *GmHs1-1* for hard-seed in soybean^10^, and *G* for dormancy in soybean, rice and tomato^11^. However, the number of genes identified through this method is rather limited. In contrast, a bottom–up approach can identify a greater number of putative domestication loci, e.g. 55 loci in rice^12^, 484 in maize^13^ and 230 in soybean^14^. A major limitation of the bottom–up approach is that resolving the functional consequences of the region/s of interest remains a complex experimental challenge.

Here, we used a bottom–up approach to show that a selective locus containing the transcription factor RR2b in the soybean cytokinin signaling pathway is situated among several pathogen–resistance genes. RR2b is a cytokinin–pathway activator and a repressor of root nodulation. Intriguingly, RR2b controls soybean pathogen resistance by regulating generation of reactive oxygen species, and balances soybean yield and disease resistance through the fine–tuning of its transcription levels. This arises from differential repeats of an ATT insertion element in the *RR2b* promoter region that alters its expression levels.

## Results

### *RR2b* was strongly selected during soybean domestication for its transcriptional activity

Because cytokinin has major impacts on many important traits in soybean, including seed size, nutrient use, root architecture and root nodulation, we assessed whether components of the cytokinin pathway (Supplementary Table 1) were selected during soybean domestication and subsequent improvement breeding, by plotting nucleotide diversity (π) and divergence index (*F*_st_). Among all 147 annotated genes involved in cytokinin biosynthesis, transportation and signal transduction in soybean, only one, *RR2b* (*Glyma.15G145200*), which encodes a type-B response regulator transcription factor, is located in a genomic region subjected to both a domestication and improvement selective sweep (Figure 1a, top). This hints at the importance of this region for agronomically desirable traits and soybean performance, and is consistent with a previous observation of genes related to domestication and improvement^14^. In this selection–sweep region, there are 10 annotated disease–resistance or drought–induced genes, implying that *RR2b* was potentially co–selected along with these resistance–related genes (Supplementary Figure 1a). *RR2b* is also located in certain reported disease–resistance–related QTLs, including for *Phytophthora sojae* (the causal pathogen of stem and root rot), *Fusarium solani* (sudden-death syndrome), *Phakopsora pachyrhizi* (soybean rust), *Heterodera glycines* (nematode), and isoflavone–metabolism genes, including genistein, glycitein and isoflavone (Supplementary Figure 1b), all of which contribute to defenses against biotic and abiotic stress. Interestingly, an InDel of varying (ATT) repeats was found in the *RR2b* promoter at the –100bp position (Figure 1a, bottom). Based on the number of these (ATT) repeats, we classified the published core–soybean germplasm population^14^ into three haplotypes: haplotype 1 (HT1) containing 15 or 16 (ATT) repeats, haplotype 2 (HT2) containing fewer than 14 (ATT) repeats, and haplotype 3 (HT3) containing 17 or more (ATT) repeats. A survey of the core soybean population comprising 296 sequenced accessions found that all three haplotypes have comparable frequencies in wild soybean *G. soja* (36%, 28%, and 36%, respectively). In landraces, the portion of HT1 increased to 79%, with HT2 and HT3 reduced to 3% and 18%, respectively. In *G. max* cultivars, HT1 frequency is predominant (90%) over HT2 (1%) and HT3 (9%) (Figure 1b). These observations affirm that the genomic region in which *RR2b* is located was selected during both soybean domestication and its subsequent improvement breeding.

**Figure 1.**
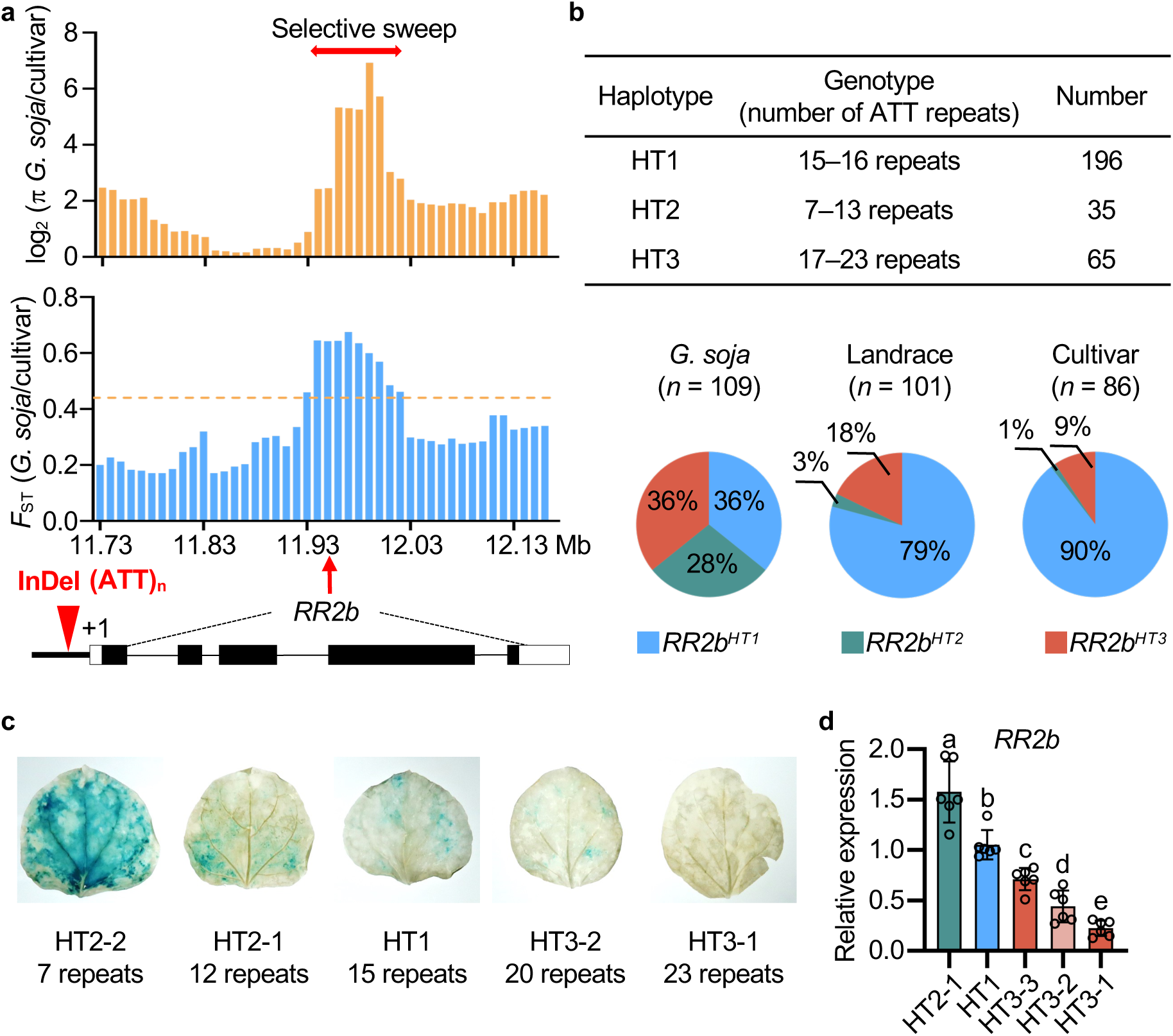
*RR2b* was selected during soybean domestication based on its transcriptional activity. (a) Nucleotide diversity (ρε, top) and population-genetic differentiation (*F*_ST_, middle) at the *RR2b* locus. The double-ended red arrow indicates the selective sweep and the dashed orange line denotes the *F*_ST_ threshold. Bottom: *RR2b* genomic structure and its position on chromosome 15 in the analyses above. (b) Distribution of three main *RR2b* haplotypes (HT) in the core collection of 296 sequenced soybean accessions. (c) Histochemical analysis of GUS expression reporting the activity of different *RR2b* promoters containing various (ATT) insertion repeats in transiently transgenic *Nicotiana benthamiana* leaves. (d) *RR2b* expression levels in the roots of different soybean haplotypes. Relative expression levels were normalized to *ELF1b*. Data are presented as means ± SD from three biological replicates. At least six accessions were analyzed for each haplotype. Different letters indicate statistically significant differences at *p* < 0.05 by one-way ANOVA with multiple comparison and Tukey’s test.

Next, we tested whether the repeat insertions in the *RR2b* promoter affect its activity. Different *RR2b* promoter haplotypes fused to a *GUS* reporter (*RR2b_pro_:GUS*) with a *REN* as an internal control were transformed into *Nicotiana benthamiana* leaves for transient expression. Activity of the *RR2b* promoter negatively correlated with the length of the insertion (Figure 1c). In soybean germplasm harboring different haplotypes, HT2 (containing short repeats in the promoter) has the highest *RR2b* expression level compared to that of HT1 and HT3 with longer repeats (Figure 1d).

In sequenced natural soybean populations, the minimal repeat of the ATT insertion is seven (Figure 1b). We generated an artificial *RR2b* promoter containing no repeats (designated ‘HT–NR’) and tested its activity using transient dual–luciferase assays, along with the natural haplotypes. This confirmed that *RR2b* promoter activity is negatively correlated with the insertion–repeat number, that is, HT–NR had the highest transcriptional activity and HT3–1 (with 23 insertion repeats, the most repeats in soybean germplasms we tested) had the lowest (Supplementary Figure 2a, b). We infer that *RR2b* was selected during soybean domestication based on its differential transcriptional activity.

### RR2b is a transcriptional activator of the cytokinin signal–transduction pathway

*RR2b* is expressed in multiple organs in soybean (Supplementary Figure 3a), and its protein product can activate the marker genes *RR5a* and *RR9c* in the cytokinin signaling pathway (Supplementary Figure 3b, c). As a typical transcription factor in this pathway, RR2b is composed of a receiver domain with conserved DDK residues, and an activation domain (Supplementary Figure 3d). In Arabidopsis, the DDK domain of type-B response regulators contains phosphorylation sites and a nuclear-localization signal, and the ΔDDK–domain mutant can activate cytokinin signaling more efficiently^15^. Likewise, full–length soybean RR2b–GFP localized in the nucleus, whereas the single– and double–Asp deletion mutants failed to localize there and instead appeared in the cytosol and plasma membrane (Supplementary Figure 3e).

To understand the roles of RR2b in the soybean cytokinin pathway, we generated stable–transgenic plants over–expressing *RR2b* fused to the 3xFLAG tag (OE), together with mutant alleles by gene editing (*rr2b*), all in the W82 wild–type (WT) background (Supplementary Figure 3f, g). Knockout and over–expression of *RR2b* resulted taller and shorter seedling heights, respectively (Figure 2a, b). Both *rr2b* mutant alleles have more root nodules, and *RR2b* OE plants have fewer nodules compared to WT controls, all with comparable root architecture (Figure 2c, d). We tested the physiological responses of these genotypes to exogenous supplementation with the cytokinin analog BAP (6–benzylaminopurine). Less than 100 nM BAP did not inhibit primary root length of WT plants. Along with higher BAP concentrations, WT root length reduced, the *rr2b*–*1* mutant showed reduced sensitivity and *RR2b* OE plants were hypersensitive (Figure 2e). This supports that *RR2b* is a positive regulator of the cytokinin pathway in soybean, akin to the Arabidopsis homolog. Interestingly, root nodulation was more sensitive than root elongation to exogenous cytokinin. WT nodule number increased at 1 nM BAP and decreased at 10 and 100 nM BAP. *rr2b*–*1* nodulation was suppressed at all concentrations and *RR2b* OE plants had more nodules at 10 and 100 nM BAP (Figure 2f). These results suggested that RR2b is a negative regulator of soybean nodulation.

**Figure 2.**
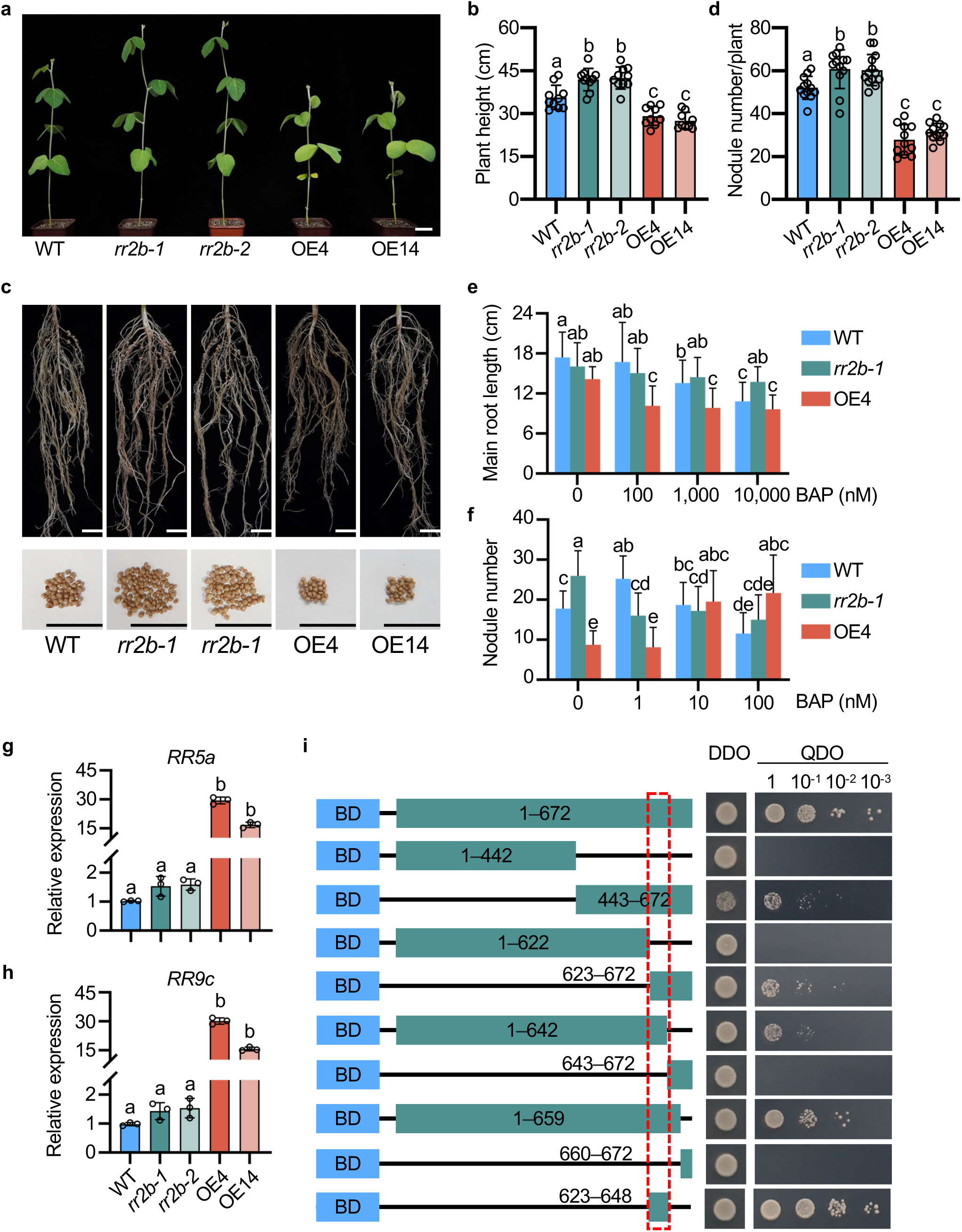
*RR2b* is a positive transcriptional regulator of the cytokinin– signaling pathway. (a) Representative phenotypes of 21–d–old wild–type c.v. W82, *rr2b* knockout and *RR2b* over–expression (OE) plants. (b) Plant height for 21–d–old W82, *rr2b* knockout and *RR2b* OE plants. Data are the means ± SD (n ≥ 15 individual plants). Three independent experiments were performed with similar results. (c, d) Nodule phenotype (c) and nodule numbers per plant (d) for W82, *rr2b* knockout and *RR2b* OE lines. Data are means ± SD (n = 18 individual plants). Three independent experiments were performed with similar results. (e) Primary root length for W82, *rr2b* knockout and *RR2b* OE transgenic lines in response to varying concentrations of exogenous BAP or mock treatment for 15 d. (f) Root-nodule numbers per plant for W82, *rr2* knockout and *RR2* OE transgenic lines in response to varying concentrations of exogenous BAP or mock treatment for 15 d. Data are means ± SD (n ≥ 13 individual plants). Three independent experiments were repeated with similar results. (g, h) Relative expression of *RR5a* (g) and *RR9c* (h) in W82, *rr2b* knockout and *RR2b* OE transgenic lines. Data are means ± SD from three biological replicates. (i) Transactivation activity of RR2b domains in yeast. Full–length RR2b or the indicated truncations (in green) were fused in–frame with the GAL4 DNA– binding domain (BD, in blue) in the pGBKT7 vector, respectively. Colony growth in selective medium indicates the transactivation of RR2b. DDO, SD/-Trp/-Leu; QDO, SD/-Trp/-Leu/-His/-Ade. The dashed–red rectangle indicates the region comprising amino acids 623–648 is sufficient and necessary for RR2b activity. Expression levels in g and h were normalized to *ELF1b*. In b, d–h, different letters indicate statistically significant differences at *p* < 0.05, as determined by multiple-comparison testing by one-way ANOVA analysis with Tukey’s test. Scale bars = 2 cm in a and c.

We also examined the induction of cytokinin-pathway marker genes (Figure 2g, h). *RR5a* and *RR9c* are dramatically induced in *RR2b* OE plants, but are comparable to WT in *rr2b*–*1* and *rr2b*–*2* mutants, probably due to the compensation or severe redundancy amongst the 24 type-A *RRs* in soybean (Supplementary Table 1). To identify the RR2b domain/s required for its activity, the functionality of truncation mutants was assayed in yeast. The RR2b C– terminal is more important for its transcription activity than the N–terminal, consistent with that in Arabidopsis. Only 26 amino acids (residues 623–648) are sufficient and necessary for this transcription activity in yeast (Figure 2i). Together, these data indicated that RR2b is a transcriptional activator of cytokinin signaling and negatively regulates nodulation in soybean.

### RR2b acts as a transcriptional repressor of soybean disease–resistance pathways to dampen ROS bursts

To further study the biological roles of the transcription factor RR2b, we performed a CUT & Tag with *RR2b–3xFLAG* transgenic plants and identified a core motif (Figure 3a). This RR2b binding motif contains the sequence TTGGAAT, which is not identical to the classic GARP-motif (G/A)GAT(T/C) of type-B RRs^15^, indicating RR2b might have distinctive function in soybean. We analyzed transcriptomes of WT, *rr2b* knockout and over–expression plants by RNA–seq. Differentially expressed genes (DEGs) between these genotypes are highly enriched in pathogen resistance–related pathways, including cell wall biogenesis, phenylpropanoid biosynthesis, plant–pathogen interaction, and flavonoid biosynthesis pathway, which are also involved in legume–rhizobia interactions for root nodulation (Supplementary Figure 4a, b). Candidate RR2b–3xFLAG target genes were filtered out by overlapping putative target sequences identified by CUT & Tag with the DEGs identified in *RR2b* OE14 plants versus WT. From this, we identified 274 candidates (Supplementary Table 2) and focused on the pathogen–related genes. Among these, known pathogen resistance genes are found, such as *PPR40* (*PENTATRICOPEPTIDE REPEAT 40*)^16^, *ATKH* (*K HOMOLOGY DOMAIN-CONTAINING PROTEIN*)^17^, and *LRR4* (*LEUCINE RICH REPEAT PROTEIN*)^18^, all of which are up– regulated in *rr2b* mutants and down–regulated in *RR2b* OE plants (Figure 3c–e), suggesting that RR2b is a transcriptional repressor of defense genes. We validated CUT & Tag peaks (Figure 3f–h) by electrophoretic mobility shift assay (EMSA). RR2b–GST directly binds to TTGGAAT *cis*-elements located in the promoters of *PPR40*, *ATKH8* and *LRR4* (Figure 3i–k). This was validated *in planta* by dual–luciferase reporter assays in which the activity of these promoters was reduced in the presence of RR2b, compared to empty–vector controls (Figure 3l–3n). All together, these results indicated that the transcription factor RR2b acts as a repressor of gene expression in these soybean disease resistance pathways.

**Figure 3.**
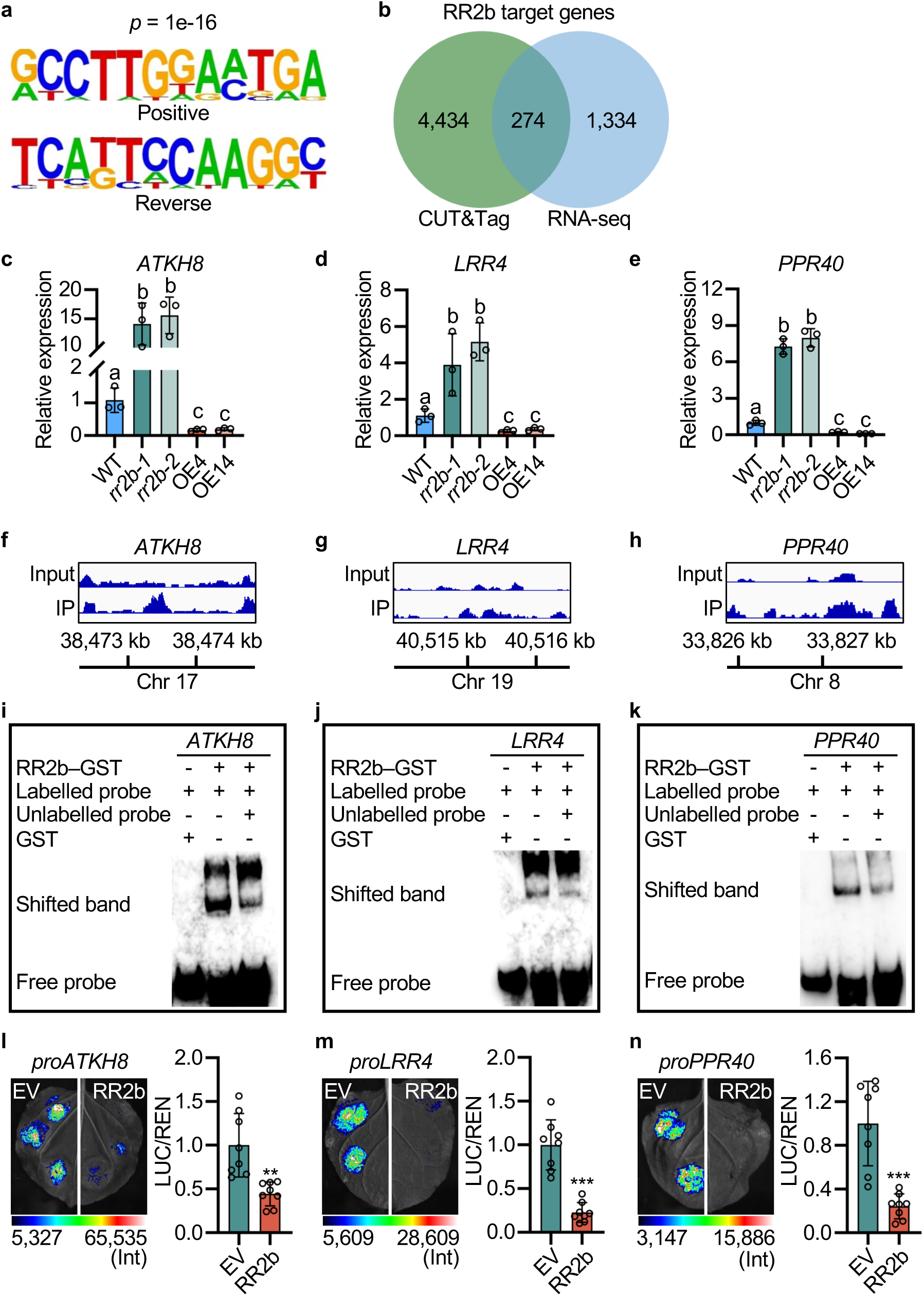
RR2b acts as a transcriptional repressor of soybean disease resistance. (a) HOMER motif analysis of regions enriched within RR2b binding regions experimentally determined by CUT & Tag. (b) Venn diagram of overlap between putative RR2b target genes identified by CUT & Tag and differentially expressed genes in *RR2b* OE14 relative to W82 as identified by RNA–seq. (c–e) Relative expression of *ATKH8* (c), *LRR4* (d) and *PPR40* (e) in W82, *rr2b* knockout and *RR2b* OE transgenic lines. Expression levels were normalized to *ELF1b* and data are means ± SD from three biological replicates. Different letters indicate statistically significant differences at *p* < 0.05 by one-way ANOVA analysis with Tukey’s test. (f–h) CUT & Tag analysis of RR2b–3xFLAG preferentially binding to the promoter of *ATKH8* (f), *LRR4* (g) and *PPR40* (h). (i–k) EMSA of GST– RR2b binding *in vitro* to *cis*-elements within promoters of *ATKH8* (i), *LRR4* (j) and *PPR40* (k). Three independent replicates were performed and a representative result is shown. (l–n) Transient dual–luciferase assays in *N. benthamiana* leaves of RR2b binding to the promoter of *ATKH8* (l), *LRR4* (m) and *PPR40* (n). Shown are relative ratios of the transcriptional activities conferred by RR2b expression to the empty–vector control. LUC/REN, ratio of firefly luciferase to *Renilla* luciferase activity. Data are presented as means ± SD (n = 8). Three independent experiments were repeated with similar results. Asterisks indicate statistically significant differences relative to the empty– vector (EV) control. Two–sided Student’s *t*–test, \*\**p* < 0.01, \*\*\**p* < 0.001.

Because RR2b binds to the promoters of select disease resistance pathway genes to inhibit their expression, we assessed disease symptoms of WT, *rr2b* knockout and *RR2b* OE plants. *rr2b* mutants show enhanced resistance to *Pseudomonas syringae pv. glycinea*, which causes soybean bacterial blight, and *RR2b* OE plants are hypersusceptible to this pathogen relative to WT (Figure 4a, b). Consistent with this, expression of pathogen resistance marker genes *PR2* and *PR5* are significantly induced in *rr2b* mutants and repressed in *RR2b* OE plants (Figure 4c, d). Production of flg22–triggered reactive oxygen species (ROS), an early hallmark of the pattern–triggered immunity (PTI) response, was dramatically higher in *rr2b*–*1* than in WT, and the ROS burst in *RR2b* OE plants was reduced compared to WT (Figure 4e, f). We then examined the relationship between RR2b and RESPIRATORY BURST OXIDASE HOMOLOG (RBOH) enzymes, the major ROS–generating system active during early stages of symbiosis and PTI responses. *RbohA*, *RbohC* and *RbohD* promoter activities were suppressed by RR2b in dual–luciferase reporter assays (Figure 4g–i), and RR2b–GST could bind *in vitro* to sequences in these promoters in EMSAs (Figure 4j–l). Furthermore, *RbohA*, *RbohC* and *RbohD* expression levels are greatly increased in *rr2b* mutants and decreased in *RR2b* OE plants compared to WT (Figure 4m–o). These functional and genetic experiments indicate that RR2b represses expression of these *Rboh* genes at the transcriptional level, thereby inhibiting soybean resistance by dampening ROS bursts.

**Figure 4.**
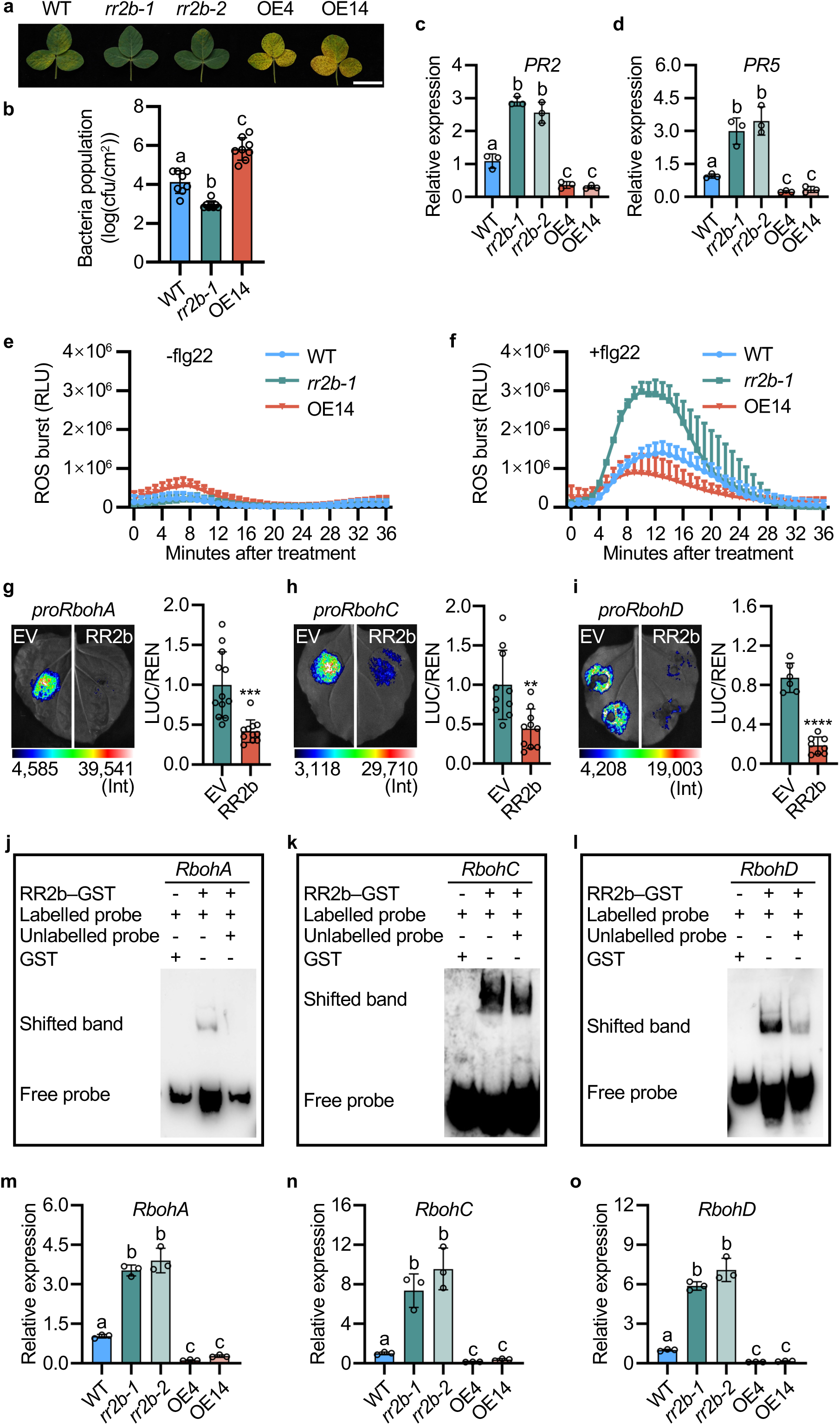
RR2b inhibits soybean disease resistance by dampening ROS bursts. (a, b) Representative images (a) and bacterial population (b) in leaves of W82, *rr2b* knockout and *RR2b* OE lines inoculated with *Pseudomonas syringae* pv. *glycinea* (*Psg*). (a) Seven days after inoculation. Scale bars = 2 cm. (b) Data are means ± SD of *n* = 8. Three independent experiments were repeated with similar results. Different letters indicate statistically significant differences at *p* < 0.05 by one-way ANOVA analysis with Tukey’s test. (c, d) Relative expression of *PR2* (c) and *PR5* (d) in W82, *rr2b* knockout and *RR2b* OE transgenic lines. Data are means ± SD from three biological replicates. (e, f) ROS production induced by mock treatment (e) or flg22 treatment (f) in 4–d–old seedlings of W82, *rr2b* knockout and *RR2b* OE lines. (g–i) Transient dual–luciferase assays of RR2b binding to the promoter of *RbohA* (g), *RbohC* (h), *RbohD* (i). Shown are relative ratios of the transcriptional activities conferred by RR2b expression to the empty–vector control. LUC/REN, ratio of firefly luciferase to *Renilla* luciferase activity. Data are means ± SD (n = 8). Three independent experiments were repeated with similar results. Asterisks indicate statistically significant differences relative to the EV control. Two–sided Student’s *t*–test, \*\**p* < 0.01, \*\*\**p* < 0.001, \*\*\*\**p* < 0.0001. (j–l) EMSA of GST–RR2b binding *in vitro* to *cis*-elements in promoters of *RbohA* (j), *RbohC* (k) and *RbohD* (l). At least three independent replicates are performed for each experiment and a representative result is shown. (m–o) Relative expression of *RbohA* (m)*, RbohC* (n) and *RbohD* (o) in W82, *rr2b* knockout and *RR2b* OE lines. Expression levels in panels c, d, m–o were normalized to *ELF1b* and data are presented as means ± SD from three biological replicates. In b–d and m–o, different letters indicate statistically significant differences at *p* < 0.05 by one-way ANOVA analysis with Tukey’s test.

### RR2b balances soybean yield and pathogen resistance

As *RR2b* was selected during soybean domestication and improvement breeding, we examined the effects of *RR2b* haplotypes on agronomic traits under natural conditions in the field at three locations spanning from 37°26’ to 39°54’ of north latitude. *rr2b* mutants were taller than WT and *RR2b* OE plants were shorter (Figure 5a, Supplemental Figure 5a*–*c). By contrast, branch number (Supplemental Figure 5d*–*f), total grain number per seedling (Supplemental Figure 5g*–*i), pod number per plant, and the proportion of the pods containing one to four seeds (Supplemental Figure 5j*–*l), are generally comparable across genotypes. Notably, one of the most important soybean domestication traits, seed size, as expressed by 100*–*seed weight, was much smaller in *rr2b* knockout plants than WT, with *RR2b* OE plants consistently producing larger seeds across all seasons and locations (Supplemental Figure 6a*–*e). These results suggested that *RR2b* positively regulates soybean seed size. Indeed, seed size positively correlates with *RR2b* transcript level in multiple field trials, with the high expression–level *RR2b* germplasm containing fewer (ATT) insertion repeats having bigger seeds than the low expression– level *RR2b* germplasm (Supplementary Figure 6f–j).

**Figure 5.**
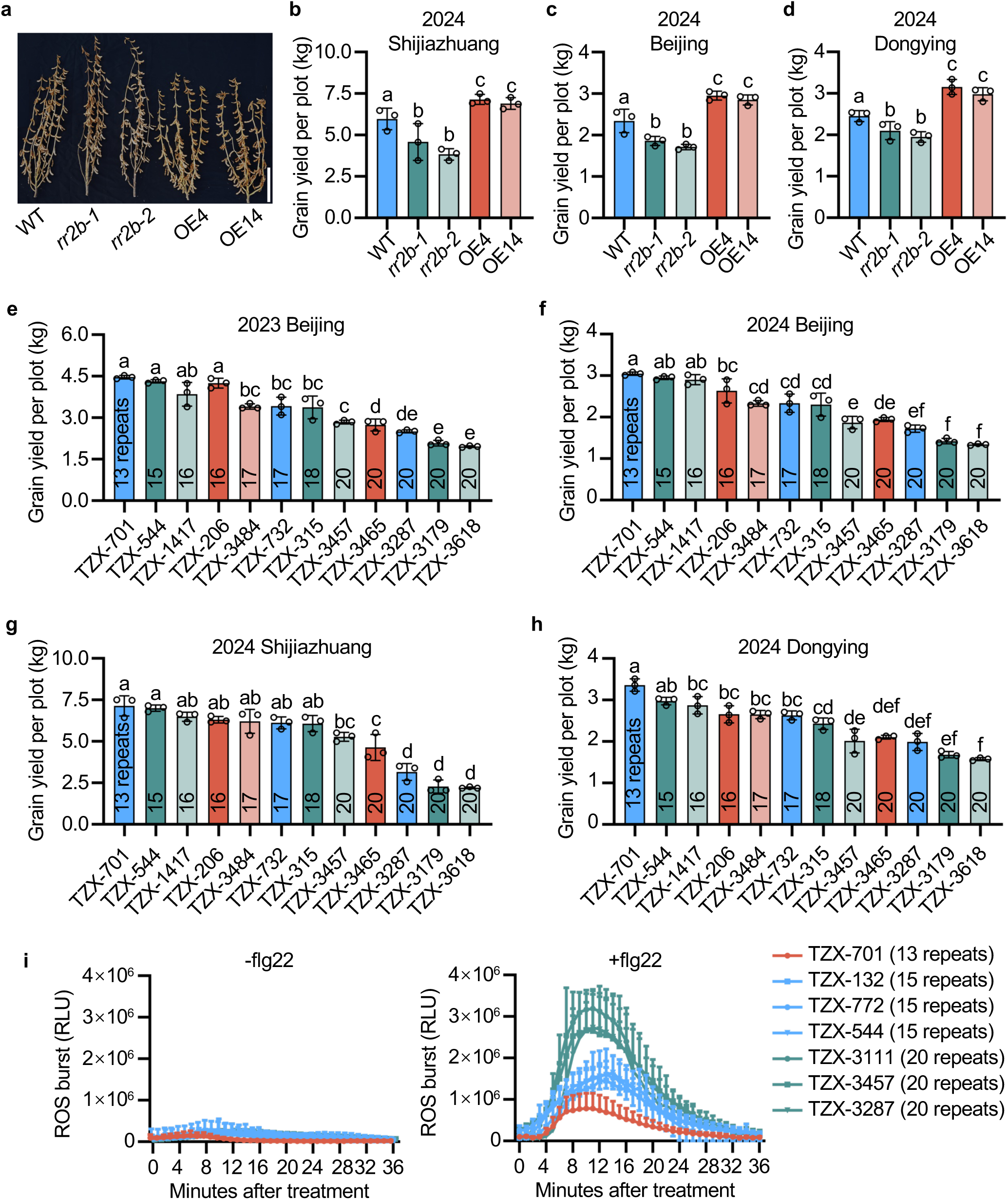
RR2b balances soybean yield and disease resistance. (a) Phenotype of W82, *rr2b* knockout and *RR2b* OE shoots at the harvest stage. Scale bar = 10 cm. (b–d) Quantification of grain yield per plot for W82, *rr2b* knockout and *RR2b* OE lines grown at Shijiazhuang (b), Beijing (c) and Dongying (d) field sites in 2024. (e–h) Quantification of grain yield per plot for soybean germplasms containing various insertion copies in the *RR2b* promoter grown at multiple field sites in 2023 and 2024. Three plots were grown for each line, with plot sizes in Shijiazhuang and Dongying of 15 m^2^ and in Beijing of 9 m^2^. (i) ROS production in 4–d–old HT1, HT2 and HT3 seedlings under mock treatment (left) or treated with flg22 (right). Data in (b–h) are means ± SD and different letters indicate statistically significant differences at *p* < 0.05 determined by one-way ANOVA analysis with Tukey’s test. For e–h, the denoted repeats refer to the copy numbers of (ATT) insertions in the *RR2b* promoter.

To place this correlation in a plant–performance context, we examined plot yield in these field conditions. *rr2b* mutants had lower yield and *RR2b* OE plants had higher yield than WT in 2024 (Figure 5b*–*d). Moreover, at these locations in 2023 and 2024, different germplasm with various insertion repeats in the *RR2b* promoter (with different expression levels) produced similar results, with the germplasm bearing shorter insertions having higher yield, and *vice versa* (Figure 5e–h). For responses to pathogen challenge, the more (ATT) repeats in the *RR2b* promoter, the stronger the ROS bursts (Figure 5i). These results collectively suggest that *RR2b* transcript level is positively related with soybean seed size and grain yield, and negatively related with pathogen resistance.

### The ABIG1–RR2B–bHLH63 module regulates soybean nodulation

RR2b negatively regulates soybean nodulation (Figure 2c, f), therefore, we aimed to elucidate the mechanistic basis of this. Several nodulation marker genes, including *NINa*, *NINb*, *ENOD40-1*, *NSP1a*, and *NSP2a*, were up*–* regulated in *rr2b* mutants and down*–*regulated in *RR2b* OE plants relative to WT (Supplementary Figure 7a–e). In CUT & Tag datasets, we also identified some known nodulation–pathway genes, including *CEP9* (*C-TERMINALLY ENCODED PEPTIDE 9*)^19^ and *NRLK* (*LEUCINE-RICH REPEAT PROTEIN KINASE FAMILY PROTEIN*)^20^ (Supplementary Figure 7f, g). EMSA confirmed that RR2b*–*GST could bind *in vitro* to *cis*–elements in the promoters of *CEP9* and *NRLK* (Supplementary Figure 7h and 7i) and RR2b could down–regulate *CEP9* and *NRLK* promoter activity in tobacco leaves (Supplementary Figure 7j, k). Consistent with this, *CEP9* and *NRLK* expression is higher in *rr2b* mutants and lower in *RR2b* OE plants compared to WT (Supplementary Figure 7l, m). These results support that RR2b represses soybean nodulation via these genes.

To identify the interaction protein partners of RR2b, a yeast two–hybrid assay was performed and 50 candidates were identified (Supplementary Table 3). Among them, a nuclear–localized basic helix–loop–helix transcription factor, bHLH63, garnered our interest. The direct interaction between RR2b and bHLH63b was confirmed in yeast (Figure 6a) and this was validated using bi*–* molecular fluorescence complementation (Figure 6b), luciferase– complementation imaging (Figure 6c) and co–immunoprecipitation experiments (Figure 6d). Toward understanding the functional relevance of the RR2b*–* bHLH63b interaction, *bHLH63b* was over–expressed in transgenic hairy roots, resulting in fewer nodules (Figure 6e, f). Dual–luciferase reporter assays demonstrated that bHLH63b could down–regulated *NINa* promoter activity, similar to the effect observed with RR2b (Figure 6g, h). These data indicate that RR2b and bHLH63b might form a transcriptional repressor complex that inhibits soybean nodulation.

**Figure 6.**
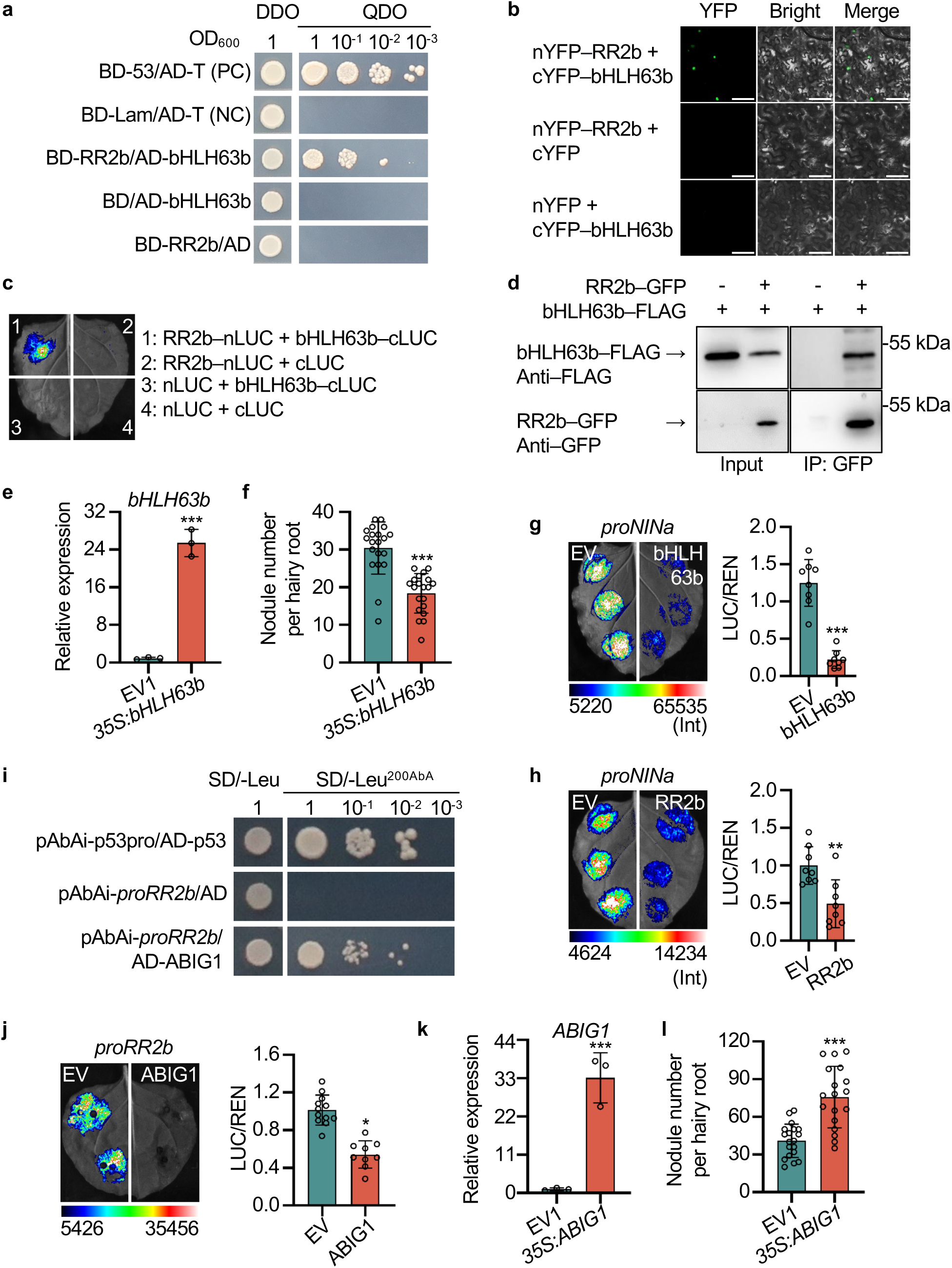
The *ABIG–RR2b–bHLH63* genetic module regulates soybean nodulation. (a) Yeast two–hybrid assay of the physical interaction between RR2b and bHLH63b. Cells were grown on DDO (Trp/Leu) or QDO (Trp/Leu/His/Ade) synthetic dropout medium and numbers at the top indicate four serial dilutions. AD, GAL4 activation domain; BD, GAL4 DNA-binding domain. (b) Bi-molecular fluorescence complementation analysis of the physical interaction between RR2b and bHLH63b in transiently transgenic *N. benthamiana* leaves. Scale bars, 100 µm. cYFP, C-terminal portion of YFP. nYFP, N-terminal portion of YFP. (c) Luciferase-complementation assay in transiently transgenic *N. benthamiana* leaves testing the interaction between RR2b and bHLH63b. Representative images of *N. benthamiana* leaves at 48 h after infiltration are shown. (d) Co-IP analysis of the physical interactions between RR2b–GFP and bHLH63b– 3xFLAG. (e) *bHLH63b* expression level in transgenic hairy roots expressing *35S_pro_:bHLH63b* or an empty-vector control (EV1). (f) Quantification of nodule number per hairy root expressing EV1 and *35S_pro_:bHLH63b*. Data are means ± SD of 20 hairy roots per construct. (g) Transient dual–luciferase assays of bHLH63b binding to the promoter of *NINa*. Shown are relative ratios of the transcriptional activities conferred by bHLH63b expression to the empty vector control. (h) Transient dual-luciferase assays of RR2b binding to the promoter of *NINa*. (i) Yeast one–hybrid assay of ABIG1 binding to the *RR2b* promoter. Numbers along the top indicate four serial dilutions. (j) Transient dual–luciferase assays of ABIG1 binding to the promoter of *RR2b*. (k) Relative *ABIG1* expression in transgenic hairy roots expressing *35S_pro_:ABIG1* or an empty– vector control (EV1). (l) Quantification of nodule number per hairy root expressing EV1 and *35S_pro_:ABIG1*. Data are means ± SD (n = 19). In c–g and h–l, three independent experiments were repeated with similar results. In g, h, j, LUC/REN, ratio of firefly luciferase to *Renilla* luciferase activity. Data are presented as means ± SD (n = 8). Three independent experiments were repeated with similar results. Asterisks indicate statistically significant differences relative to the EV control. Two–sided Student’s t–test, \*\*\**p* < 0.001, \*\**p* < 0.01, \**p* < 0.05. In e and k, gene–expression levels were normalized to *ELF1b* and data are means ± SD from three biological replicates. In panels e, f, k, l, asterisks indicate statistically significant differences relative to the EV1 control. Two–sided Student’s *t*–test, \*\*\**p* < 0.001.

Finally, we sought to identify the upstream regulatory *trans*–element(s) of the *RR2b* promoter by yeast–one hybrid (Supplementary Table 4). ABIG1 (ABA INSENSITIVE GROWTH 1), the homolog of an abiotic stress–related transcription factor ABIG1 in Arabidopsis^21^, could activate the *RR2b* promoter in yeast (Figure 6i). Dual–luciferase reporter assays showed that ABIG1 repressed the *RR2b* promoter (Figure 6j) and *ABIG1* over–expression in transgenic hairy roots results in more nodules (Figure 6k, l), like *rr2b* knockout mutants (Figure 2c, d), indicating that ABIG1 positively regulates soybean nodulation by repressing *RR2b*. In sum, our data suggested that an ABIG1– RR2B–bHLH63 module acts to regulate soybean nodulation.

## Discussion

As an essential plant growth hormone with crucial developmental functions, a broader understanding of how cytokinins regulate important agronomic traits makes this classic hormone an emerging genetic target for crop yield improvement^22–26^. Cytokinin signaling in plant cells is transduced through a two-component system, with B-type RRs serving as positive transcription factors^27^ that are involved in regulating soybean seed size and phosphorus uptake^28,29^. Here, we demonstrated that RR2b positively affects soybean grain yield by increasing seed size, while negatively regulating pathogen resistance by suppressing the ROS burst typical of PTI. This transcription factor was artificially selected during soybean domestication based on its transcript level and balances yield and pathogen resistance, making it a promising target for optimizing both soybean yield and resistance (Supplementary Figure 8).

Artificial selection of crops occurs in two stages, domestication and improvement. Early domestication steps change fundamental characteristics of the wild progenitor, resulting in landrace varieties, which provide the genetic materials for breeders to later select improved varieties and inbred lines. Subsequent diversification or improvement emerges as crops spread to different regions and undergo more conscious selection that can enhance traits that influence agricultural productivity and performance, including yield and resistance to stress^30^. In response to changing environments during the early domestication steps, breeders have selected plants that are better adapted to local conditions. This selection process has led to the emergence of landraces, which are traditional varieties chosen by farmers for their suitability to local conditions and food preferences. Although these varieties typically exhibit low yield potential, they are often characterized by genetic diversity and resilience to environmental stresses^31^. Here, a particularly intriguing phenomenon is that the high–yield *RR2b* haplotype HT2 was initially lost during soybean domestication from the wild relative to landrace varieties, followed by a decline in the prevalence of the highly disease–resistant haplotype HT3 during improvement from landraces to cultivar varieties. One possible explanation is that, in the early stages of cultivation, farmers had limited effective methods for disease control. Therefore, it is essential to first preserve varieties that are disease–resistant, followed by the selection of varieties that achieve an optimal balance between disease resistance and yield.

To date, research on the selection of disease–resistance traits during crop domestication remains fairly limited. Zhou *et al*.^14^ demonstrated that the region harboring the root–knot nematode resistance locus *Rhg1* in soybean was subject to selection. Additionally, a pan–transcriptome analysis revealed that disease–resistance genes were also selected during barley domestication^32^. The prevailing view suggests that the reduction of genetic diversity during domestication may lead to a ‘broad susceptibility’ to newly emerging herbivores and pathogen strains, thereby compromising long–term crop sustainability^33–35^. In legumes, half of the annotated resistance–related sequences in soybean and chickpea were lost by domestication and subsequent improvement^14,36^. Therefore, retrieving ancestral disease–resistance genes lost due to genetic bottlenecks during artificial selection from wild relatives or landraces, or developing disease–resistance genes preserved in cultivated varieties, such as *RR2b* identified here, presents feasible strategies for enhancing soybean disease–resistance breeding.

Both plants and animals produce ROS, which not only have a direct antimicrobial effect but also serve as signaling molecules to activate pathogen– resistance responses^37^. Exogenous application of high concentration of cytokinins or increasing the content and signaling of endogenous cytokinins through genetic means in Arabidopsis can enhance defenses against pathogens. In this process, ARR2 plays positive roles by regulating the salicylic acid pathway and ROS homeostasis, which involves apoplastic peroxidases PRX4, PRX33, PRX34, and PRX71, but not the RBOH class of NADPH oxidases^38,39^. However, our results suggest a negative role for RR2b in soybean resistance through a RBOH-dependent ROS burst (Figure 4). This discrepancy might be caused by the different mechanisms of ROS generation. Indeed, RR2b plays distinct roles in various physiological processes and signaling pathways. For instance, here, RR2b activates the cytokinin pathway in soybeans, akin to its homologous gene in Arabidopsis^15^. Meanwhile, it also functions as a transcriptional repressor, inhibiting disease–resistance pathways and root–nodule formation in soybean. Likewise, mutation of a B–type RR, *MtRRB3*, lead to fewer root nodules in *Medicago truncatula*, indicating its role as a positive regulator of the nodulation pathway^40^. In contrast, our results indicate that RR2b represses the nodulation pathway, despite its capacity to activate cytokinin signaling as a transcription factor, which aligns with a previous study in soybean^41^. This inconsistency may be attributed to species differences or variations between determinate and indeterminate nodules in Leguminosae.

Crop domestication is typically examined from evolutionary and genetic perspectives, primarily focusing on key agronomic traits referred to as domestication–related traits. However, there has been limited research on ecological domestication traits, such as growth and resource–acquisition rates, interactions with microbes, insects and pollinators. The significance of such traits lies not only in yield (i.e. plant fitness) but also includes various ecological effects, such as carbon and nitrogen sequestration, nutrient use and cycling in the soil, the ecological balance amongst other community members, as well as implications for climate change. Relatively fewer ecological domestication trait studies have focused on crop adaptation to climate change and photoperiod^42-^_45_. Here, we revealed the artificially selected transcription factor RR2b in the cytokinin pathway orchestrates domestication–related traits (i.e. yield) and ecological domestication traits (i.e. pathogen resistance) during soybean domestication. When pathogens invade agricultural systems and lead to disease, the balance between yield and disease resistance becomes important, and rapid and precise regulation at the level of transcription may represent a shortcut to achieve this balance.

The growth–defense trade–off is regarded as one of the fundamental principles of ‘plant economics’^46^. Strong trade-offs caused by gene pleiotropy are one of the biggest barriers to crop improvement. For example, the *gl4* mutation in the domestication gene in African cultivated rice *O. glaberrima* reduces seed shattering, albeit at the expense of seed size^47^. In Asian rice, *semi-dwarf1* (*sd1*) reduces lodging risk but also decreases plant biomass and nitrogen-use efficiency^48^. A gain–of–function of *IPA1* (*IDEAL PLANT ARCHITECTURE 1*) results in larger panicles in rice and enhanced disease resistance, but leads to fewer tillers^49^. In this study, pleiotropy of *RR2b* results in a trade–off between yield and disease resistance. There are two feasible approaches to decouple this pleiotropy. One is to target a gene regulatory element to modify its *cis*–regulatory regions, thereby separating the multiple functions of RR2b by altering the developmental stages and tissue–specificity of gene its expression, as performed in other crops^50^. The second entails identifying distinct protein function domains that may contribute to pleiotropy and constructing various truncated or mutated versions of this transcription factor to fulfill the necessary functions under different environmental conditions. Our identification of the minimal, sufficient and necessary sequence for RR2b transcriptional–regulation activity provides the possibility for this approach (Figure 2i). In future soybean breeding, disrupting *RR2b* pleiotropy and addressing linkage drag will offer new strategies to decouple the yield–disease resistance trade–off, ultimately leading to the development of high–yielding and disease–resistant varieties.

## Declaration of interests

This work has been filed for patent applications with inventors B.R., Q.M., M.Z., Q.W., and X.H.

## Author-contribution statement

Q.M., L.L, Z.T., and B.R. conceived the research, Q.M., X.H., Q.Y., Y.G., and Y.L. performed the experiments, Q.M., L.L., Z.T., and B.R. analyzed the data, B.R. wrote the manuscript with input from Q.M., L.L., and Z.T.

## Acknowledgements

We thank C. Tian (China Agricultural University) for the *Bradyrhizobium diazoefficiens* strain USDA110, J. Zhang (Institute of Genetics and Developmental Biology) for the pCAMBIA2300-nYFP and pCAMBIA2300-cYFP vectors, J. Zhou (Yazhouwan National Laboratory) for the pCAMBIA1300-nLUC and pCAMBIA1300-cLUC vectors, X. Li (Huazhong Agricultural University) for the pEZRK vectors, D. Zhang (Shandong Agriculture University) for some soybean germplasms, and S. Hu (Institute of Genetics and Developmental Biology) and X. Liu (Institute of Genetics and Developmental Biology) for field work.

## Funding

This work was supported by the National Key Research and Development Program of China (2021YFF1000102), the CAS Project for Young Scientists in Basic Research (YSBR-011), and STI 2030–Major Project (2023ZD04072).

## Data availability

All data are available in the National Genomics Data Center (https://ngdc.cncb.ac.cn/), Beijing Institute of Genomics, Chinese Academy of Sciences, under the BioProject number PRJCA040019.

## Materials and methods

### Plant materials and growth conditions

*Glycine max* cv. Williams 82 (W82) was used as the wild type for gene cloning, gene expression and hairy–root transformation experiments. Soybean seeds were sown in sterile vermiculite for germination. The plants were propagated in a greenhouse (16-h light/8-hour dark cycle at 25°C and 65% relative humidity) equipped with LED lights at 160 µmol m^-2^s^-1^ light intensity. After germination for 5 d, seedlings were inoculated with a suspension of *Bradyrhizobium japonicum* USDA110 (OD_600 nm_ = 0.08) and the root nodules were examined 21 d later.

### Soybean genomic DNA extraction

DNA was extracted from the leaves using a CTAB (cetyltrimethylammonium bromide)–based method. Leaves were ground with 600 µL of CTAB solution and incubated in an oven at 65°C for 20 min. Three hundred microliters of chloroform was added and the mixture was shaken vigorously before centrifugation at 13,000 ×*g* for 10 min. Subsequently, 350 µL of the supernatant was aspirated and 700 µL of anhydrous ethanol was added, mixed well, and then transferred to –20°C for 30 min. A 13,000 ×*g* centrifugation was then performed for 10 min. The supernatant was discarded and the sediment was washed in 70% v/v ethanol twice. The samples were then centrifuged at 13,000 × *g* for 5 min, and the supernatant was discarded. Each sample was dissolved in 100 µL of water after desiccation.

### RNA extraction and quantitative PCR analysis

Total plant RNAs were extracted using TRIzol Up Kit (TransGen, Beijing, China). Genomic DNAs were removed and cDNA was synthesized from 500 ng of RNA using the HiScript III RT SuperMix for qPCR with gDNA wiper (Vazyme, Nanjing, China). The qRT–PCR reaction was performed using Taq Pro Universal SYBR qPCR Master Mix (Vazyme, Nanjing, China), and the primer pairs used for qRT–PCR are shown in Supplementary Table 6. The PCR reaction system contained 5 µL of 2 × Taq Pro Universal SYBR qPCR Master Mix, 0.2 µL each of Primer F and Primer R (10 µM), 1 µL of template cDNA (50 ng/µL), and 3.6 µL of RNase-free ddH₂O. The PCR protocol consisted of an initial denaturation at 95°C for 1 min (1 cycle), followed by 40 cycles of denaturation at 95°C for 5 s and annealing/extension at 60°C for 30 s, with a final melting curve analysis. *GmELF1b* (*Glyma.02G276600*) expression was used as an internal reference^51^. Fold changes were calculated from the 2^−ΔΔCt^ values.

### Plasmid construction

Construction of *GUS* reporter plasmids was done using genomic DNA obtained above as a template, and PCR reactions were performed using primer pairs to obtain a series of promoter sequences (∼3,000 bp) carrying different *RR2b* haplotypes. The promoter sequences were cloned into the intermediate vector pGWCm through the In-Fusion reaction system (Vazyme, Nanjing, China). Using Gateway LR cloning, the intermediate vector was subjected to an LR reaction with the vector pBGWFS7 to obtain *GUS* reporter vectors.

Construction of *LUC* reporter plasmids using genomic DNA as a template, PCR reactions were performed using primer pairs to obtain a series of promoter sequences (∼3000 bp). The promoter sequences were cloned into the CP461 vector through the In-Fusion reaction system (Vazyme, Nanjing, China).

W82 cDNA was used as template for PCR using primer pairs to obtain the target coding sequences. The above coding sequences were cloned into pTF101 through the In-Fusion reaction system (Vazyme, Nanjing, China) to obtain an over–expression vector in which RR2b was fused to a triple FLAG tag (3xFLAG) at the C-terminus.

### Genetic-diversity analysis

SNPs and InDels data from previously reported were used for diversity analysis of *RR2b*^14^. The SNPs and InDels with missing data >10% or MAF<5% were filtered. Accessions were divided into three populations: *G. soja*, landraces, and cultivars. π was calculated using a 20-k-2-k sliding window. After filtering the windows with <10 SNPs/InDels in wild and 0 SNPs/InDels in cultivated populations, we calculated the ratio of diversity (ν_wild_/ν_cultivated_) for each window. *F*_ST_ values were calculated with a 20-k-2-k sliding window using VCF tools^52^ to calculate the pairwise genomic differentiation for wild and cultivated populations of soybean. The top 5% genome sequences were determined as selective sweeps.

### Haplotype analysis of *RR2b* in the soybean population

SNPs and InDels analyses of the 3–kb promoter region and full–length coding sequences of *RR2b* were done using 296 sequenced varieties in a natural population^53^. The SNPs and the InDels were filtered by applying a MAF >5% cutoff, missing rate <10%, synonymous SNV and non–functional SNP mutation^54^.

### Hairy–root transformation and generation of stable over–expression and gene–edited soybeans

The over–expression plasmids were transformed into *Agrobacterium rhizogenes* K599, which was used to infect soybean seedlings following a previously published protocol^55^.

For CRISPR–Cas9 vector construction, the pYLCRISPR/Cas9-DB vector was used as previously reported^56^ and small guide RNAs were designed using CRISPR-P2.0 software (http://crispr.hzau.edu.cn/CRISPR2/)^57^. Two highly specific sequences in the exonic region of the *RR2b* gene, TCGAGTTCTCCTCTGAAAGCCGG and CTTGCCTTATGATCCTTGAGAGG, were selected as potential targets for gene editing. The *RR2b* knock–out vector was transformed into *A. tumefaciens* EHA105. *Agrobacterium tumefaciens*– mediated genetic transformation was used to obtain T_1_-generation *RR2b* knockout lines. In order to analyze whether *RR2b* was edited and its editing effect, its integrity in the T_1_ generation was detected by DNA sequencing. T_2_ and T_3_ Cas9-free homozygotes were analyzed for phenotypes, cytokinin responses, and agronomic traits.

For generating over–expression stable transgenic plants, W82 cDNA was used as a template for PCR to obtain the *RR2b* coding sequence for cloning into the pWMV078 vector through the In-Fusion reaction system. The vector was transformed into *A. tumefaciens* EHA105 and genetic transformation of soybean was performed based on a previously protocol^58,59^. Putative transgenic plants were genotyped by PCR and lines with at least 5-fold higher expression than that of the wild type were deemed to be over–expression plants.

### Analysis of *RR2b* promoter haplotype transcriptional activity

PCR was performed to obtain a series of promoter sequences (∼3,000 bp) carrying different *RR2b* haplotypes. The promoter sequences were cloned into the intermediate vector pGWCm through the In-Fusion reaction system (Vazyme, Nanjing, China). Using Gateway LR cloning, the intermediate vector was subjected to an LR reaction with the vector pBGWFS7 to obtain *GUS* reporter vectors. The above vectors were transformed into *A. tumefaciens* GV3101, mixed in equal proportions with a strain harboring the P19 suppressor of silencing, and uniformly infiltrated into the lower epidermis of *Nicotiana benthamiana* leaves, and maintained for 48 h to obtain transiently transgenic leaves. GUS expression was analyzed by histological staining^58^.

To verify the transcriptional activity of the different haplotypes, genomic DNA was used as template for PCR promoters carrying the different *RR2b* haplotypes. The above sequences were cloned into CP461 vector through the In-Fusion reaction system (Vazyme, Nanjing, China), obtaining the dual-luciferase reporter plasmid *RR2b^HT^ :LUC*. The above vectors were transformed into *A. tumefaciens* GV3101, mixed with P19 in equal proportions, and uniformly infiltrated into the lower epidermis of *N. benthamiana* leaves, and maintained for 48 h to obtain transiently transgenic leaves. Dual luciferase assay reagent (Beyotime, Shanghai, China) was used for luciferase imaging using the sea kidney luciferase gene as an internal control.

### Yeast two–hybrid (Y2H) assay

For testing auto–activation in yeast, the full–length coding sequence of *RR2b* and different–length coding sequences were obtained by PCR. By homologous recombination, PCR products were ligated into pGBKTT7 vector and different lengths of BD vector and AD vector were co–transformed into Y2H Gold strain and grown on SD/-Trp/-Leu (DDO) medium. Interactions were assayed by spreading 3 µL of suspended transformed yeast on plates containing SD/-Trp/-Leu/-Ade/-His (QDO) medium. The interactions were observed after 3–4 d of incubation at 30°C.

To identify RR2b–interacting proteins, we performed a Y2H screening using the GAL4 yeast two–hybrid system (Coolaber, Beijing, China). RNA was extracted from all tissues of W82 and used to make cDNA for the Y2H library, which was constructed by OE Biotech (Shanghai, China). The Y2H strain containing the BD vector with the full–length coding sequence of *RR2b* was grown on SD/-Trp/-Leu (DDO) medium in a low–rate co–culture with the AD library strain. Clones on DDO were spread 3 µL of suspended transformed yeast on plates containing SD/-Trp/-Leu/-Ade/-His (QDO) medium. Clones on QDO were sequenced and compared on the SoyBase to obtain the candidate interacting proteins.

The full–length coding sequence of *RR2b* was amplified by PCR, and cloned into the pGBKT7 (bait) vector to generate the BD–RR2b plasmid. Full-length coding sequence of *bHLH63b* was amplified and inserted into pGADT7 (prey) vector yielding plasmid AD–bHLH63b. Bait plasmids were linearized and transformed into Y2H Gold. Positive cells were then transformed with the AD– bHLH63b plasmid. Transformation was confirmed by growth on SD/-Trp/-Leu (DDO) medium. Interactions were assayed by spreading 3 µL of suspended transformed yeast on plates containing SD/-Trp/-Leu/-Ade/-His (QDO) medium. The interactions were observed after 3–4 d of incubation at 30 °C. Primers used are listed in Supplementary Table 6.

### Bimolecular fluorescence–complementation (BiFC) assay

The coding sequences of the target proteins were obtained using cDNA template and cloned into the pCambia2300-NYFP and pCambia2300-CYFP vectors through the In-Fusion reaction system. The vectors were transformed into *A. tumefaciens* GV3101, mixed with P19 in equal proportions, and uniformly infiltrated into the lower epidermis of *N. benthamiana* leaves, and incubated for 48 h to obtain transgenic leaves. Leaves were imaged using a ZEISS LSM 980 super-resolution confocal microscope.

### Luciferase complementation assay

*RR2b* was cloned in–frame and upstream of the sequence encoding the N-terminal half of firefly luciferase in pCAMBIA1300-NLUC, while *bHLH63b* was cloned in-frame and downstream of the C-terminal half of luciferase in pCAMBIA1300-CLUC. The resulting vectors, as well as empty plasmids, were introduced into *A. tumefaciens* GV3101, and transiently expressed in *N. benthamiana* leaves as described above. Luciferase activity from *N. benthamiana* leaves was detected 2 d after infiltration with a chemiluminescence imager with a cooled CCD camera.

### Co-IP assay

The RR2b–GFP vector and bHLH63b–3XFLAG vector were co–expressed in *N. benthamiana* leaves, and total proteins were extracted in buffer (50 mM HEPEs, pH 7.5, 150 mM, NaCl, 10 mM EDTA, pH 8.0, 1% v/v Trion X-100, 10% w/v sucrose, 2 mM DTT, 1 mM PMSF, 25 µM MG132, 1× protease inhibitor cocktail) and incubated with 20 µL anti–GFP beads (AlpaLifeBio, Guangdong, China) for 2 h at 4°C. Beads were washed with buffer three times, to which 50 µL of 1× SDS–PAGE sampling loading buffer was added and heated at 95°C for 10 min. Proteins co–precipitated and extracted from plant tissues were separated by 10% SDS-PAGE and transferred to PVDF membranes. After blocking with 5% non–fat milk in TBST for 1 h at room temperature, membranes were incubated overnight at 4°C with primary antibodies (diluted 1:3,000 in blocking buffer), followed by HRP–conjugated secondary antibodies (1:10,000 dilution, 1 h at room temperature). Signals were detected using an ECL substrate and visualized with a chemiluminescence imaging system^60^.

### Yeast one–hybrid (Y1H) assays

A 500–bp fragment of the *RR2b* promoter was amplified by PCR, and cloned into the pAbAi (bait) vector to generate a pAbAi-*_pro_RR2b* plasmid. The full– length *ABIG1* coding sequence was amplified and inserted into pGADT7 (prey) vector yielding plasmid AD–ABIG1. The bait plasmids were linearized and transformed into the yeast strain Y1H Gold. Positive cells were then transformed with AD–ABIG1 plasmid. The DNA–protein interaction was determined based on the growth ability of the co–transformants on SD/-Leu medium with 200 ng/mL Aureobasidin A (AbA) (CAS 127785-64-2, Yeasen).

### CUT & Tag–seq

CUT&Tag was performed using Hieff NGS® G-Type In-Situ DNA Binding Profiling Library Prep Kit for Illumina (Yeasen, Shanghai, China) according to the manufacturer’s protocol and was performed as described previously^61^. Wild–type W82 and *RR2b*–*3_X_FLAG* OE plants were grown in vermiculite for 5 d and then inoculated with *B. japonicum* USDA110 (OD_600 nm_ = 0.08), and fresh root tissue was collected after 3 d of continued growth. The nuclei of soybean root tissues were extracted and incubated with primary and secondary antibody at room temperature for 2 h and 0.5–1 h, respectively. The samples were incubated at 37°C for 1 h for transposase activation, further digested with proteinase K, and DNA was recovered by magnetic beads. The DNA library was amplified and purified, and sent to NovaSeq for sequencing at Novogene (Beijing, China).

### RNA–seq sample preparation and sequencing

Roots of WT, *rr2b* knockout and *RR2b–3xFLAG* OE plants were collected for RNA–seq analysis. Three biological replicates were performed for each sample. The Illumina HiSeq 2000 platform was used to generate 150–bp paired–end reads. Detailed bioinformatic analyses were performed as previously described^54^. Briefly, the high–quality sequencing reads were mapped to the reference genome with Hisat (v.2.2.1) and gene–expression counts were calculated using StringTie (v.1.3.4d). Differential gene–expression analysis was done using R–edgeR library (https://bioconductor.org/packages/release/bioc/html/edgeR.html).

### Transient dual–luciferase assay

To validate RR2b target genes, promoter sequences of soybean *NINa*, *CEP9*, *NRLK, PPR40*, *RbohA*, *RbohC*, *RbohD, RR5a, RR9c*, *ATKH8* and *LRR4* genes were obtained by PCR using W82 genomic DNA. PCR products were cloned into the CP461 vector through the In-Fusion reaction system (Vazyme, Nanjing, China) to obtain dual–luciferase reporter vectors. pTF101 was used to construct the over–expression vector *35S_pro_:RR2b* for the full-length coding sequence of *RR2b*. The above vector was transferred into *A. tumefaciens* GV3101, mixed with P19 in equal proportions, and uniformly infiltrated into the lower epidermis of *N. benthamiana* leaves, and maintained for 48 h to obtain transiently transgenic leaves. Dual luciferase assay reagent (Beyotime, Shanghai, China) was used for luciferase imaging using the sea kidney luciferase gene as an internal control.

### Electrophoretic mobility shift assay (EMSA)

EMSA was performed using the LightShift Chemiluminescent EMSA Kit according to the manufacturer’s protocol and as described previously^62^. GST and GST**–**RR2b were expressed in *E. coli* Rosetta and purified. The binding activity of the protein was analyzed using an oligonucleotide probe labeled with biotin at the 5′ end (Sangon Biotech, Shanghai, China). For unlabeled probe competition, 20 and 200-fold excess unlabeled probes were added to the reactions.

### Reactive–oxygen burst measurements

ROS production assay was performed as previously described^63^. Briefly, wild– type W82, *rr2b* knockout and *RR2b–3xFLAG* OE plants were grown in vermiculite for 4 d and fresh root tissue was cut into pieces about 3 mm long and transferred to a 96–well plate with 200 µL of ddH_2_O. Incubation in low light overnight allowed the reactive oxygen species produced by the tissue damage to quench themselves. The next day, the water was discarded and 200 µL of a buffer solution containing 20 µM luminol and 10 µg/mL peroxidase with the appropriate concentration of elicitors such as 100 nM flg22 was added. Detection of fluorescent signals was performed using an EnVision plate reader (Perkin Elmer).

### Disease–resistance testing

Soybean plants were grown in vermiculite for 14 d and the first ternately compound leave were selected for inoculation. Inoculation with *Pseudomonas syringae* pv. *glycinea* (OD_600 nm_ = 0.05) was performed on both leaf surfaces and incubated at high humidity for 2 d. Bacteria in leaves were extracted with ddH_2_O and 20 µL of suspension was spread on TSA medium, then the number of colonies were counted 2 d later after growth at 28°C.

### Trait measurement and plot field tests

W82, *rr2b* knockout and *RR2b–3xFLAG* OE plants were grown in fields in Beijing, Shijiazhuang and Dongying (China) under natural conditions in 2023 and 2024. Soybean germplasm was also grown in the same fields. Each plot had three replicates of the same size, six rows were planted in each plot and plants were collected from the middle four rows for yield testing. The length of each plot was 3 m (Beijing) or 5m (Shijiazhuang and Dongying) and the spacing between rows was 50 cm. Three weeks after sowing, the seedlings were manually thinned to achieve a plant density of 1,350 plants per 100 m^2^. Plant height was measured at maturity stage. Fully filled grains were used for measuring 100–seed weight.

## Statistical analysis

Histograms were made using GraphPad Prism 8.2.1, data are plotted as means ± SD as indicated, and every dot represents each sample. A one–way analysis of variance (ANOVA) was conducted, followed by the Tukey multiple– comparisons test to compare the means of each column with the means of all other columns. Statistical significance between a treatment with a control was analyzed by two–tailed, unpaired Student’s *t*–test. Categorical data associations were evaluated using Fisher’s exact test, with exact *P* values calculated based on hypergeometric distributions.

## Supplemental figure legends

**Figure S1.**
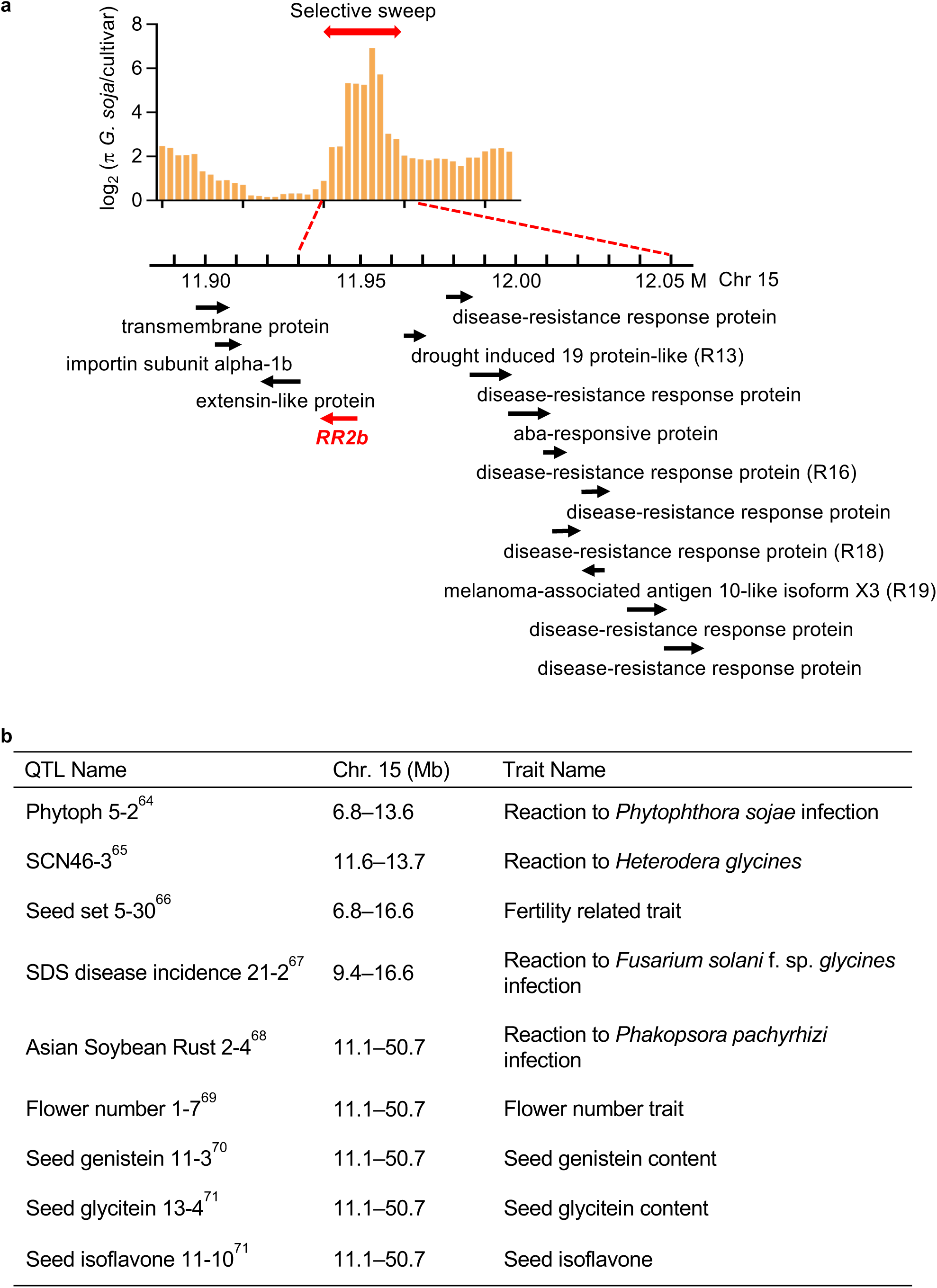
Characterization of the *RR2b* locus in the soybean genome. (a) *RR2b* is co-selected with clustered resistance genes during soybean domestication. Nucleotide diversity (ρε) at the *RR2b* locus is displayed with colored lines. Black arrows indicate the gene direction. (b) *RR2b* is situated in several reported resistance/yield-related QTLs. References are superscript on the QTL names.

**Figure S2.**
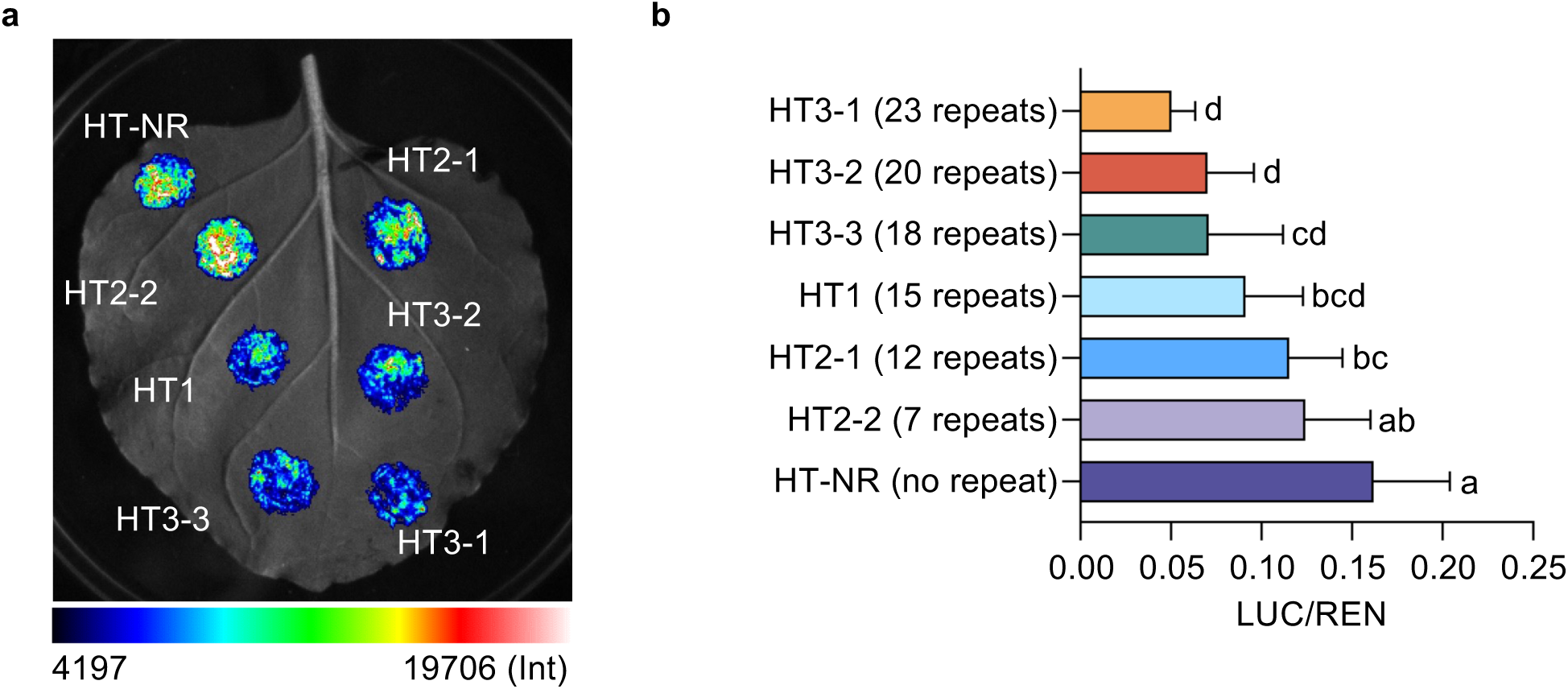
The number of insertions in the *RR2b* promoter dictates its activity. (a) Chemiluminescent image of transient dual–luciferase assays of *RR2b* promoter activity from various haplotype groups. (b) Relative LUC/REN of promoters assayed in panel a. ‘HT-NR’ (‘no repeat’) is an artificial haplotype containing zero ATT insertions in the *RR2b* promoter. Data are means ± SD (n = 8). Three independent experiments were repeated with similar results. Different letters indicate statistically significant differences in a one-way ANOVA analysis with Tukey’s test (*p* < 0.05).

**Figure S3.**
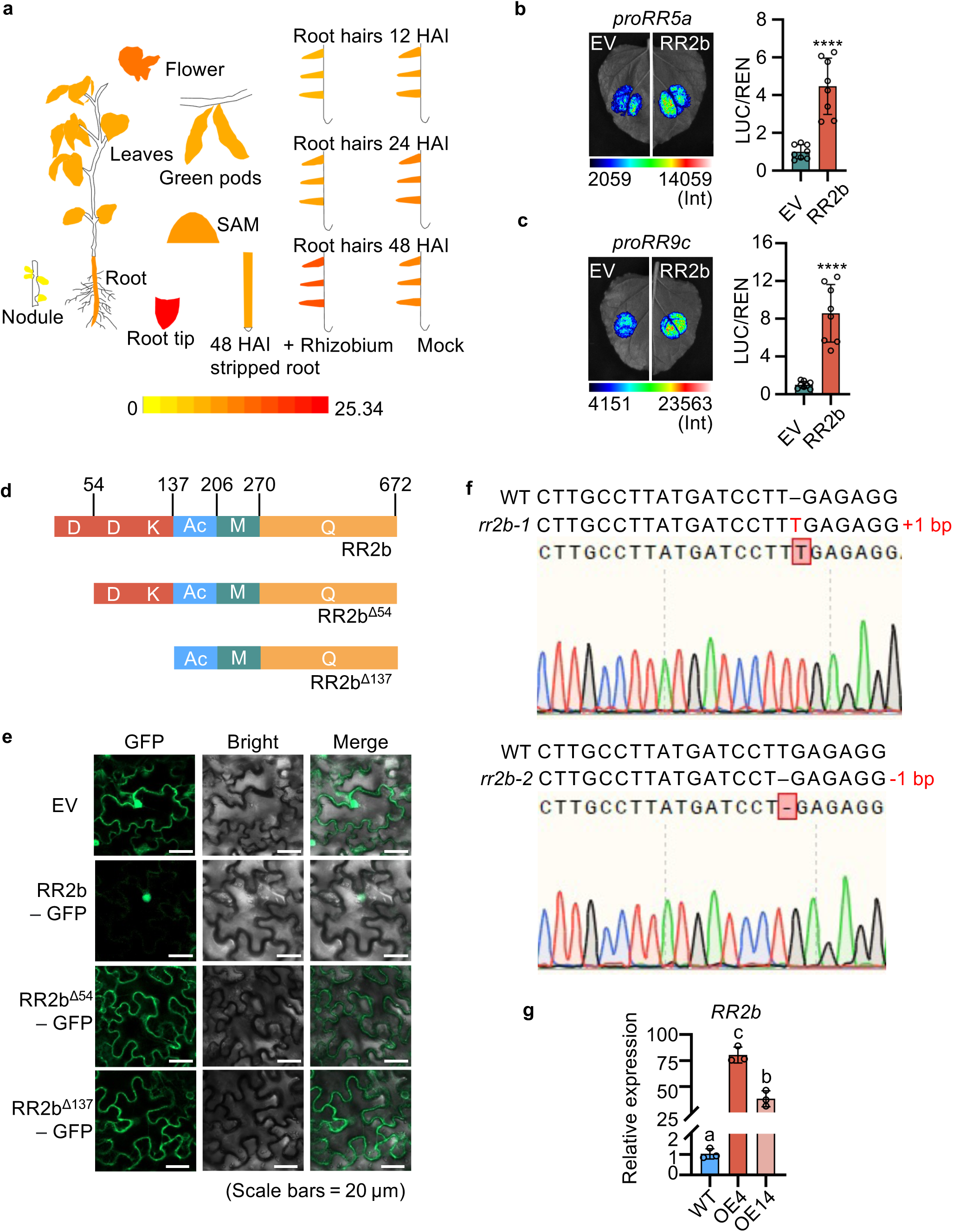
Characterization of *RR2b* tissue-level gene expression, RR2b domain structure and subcellular localization, and plant lines lacking or over–expressing *RR2b*. (a) Expression pattern of *RR2b* in various tissues obtained from the Plant eFP Viewer (https://bar.utoronto.ca/eplant_soybean/). (b, c) Transient dual– luciferase assay of RR2b activating cytokinin signaling pathway promoters *RR5* (b) and *RR9c* (c). LUC/REN, ratio of firefly luciferase to *Renilla* luciferase activity. Data are means ± SD (n = 8). Three independent experiments were repeated with similar results. Asterisks indicate statistically significant differences relative to the EV control. Two–sided Student’s *t*–test, \*\*\*\**p* < 0.0001. (d) Schematic of the structure of RR2b. RR2b is the full–length version, RR2b^Δ54^ is the first Asp-deletion version, and RR2b^Δ137^ is the double Asp-deletion version. (e) Subcellular localization of RR2b–GFP. Constructs (*35S_pro_:GFP*, *35S_pro_:RR2b*– *GFP*, *35S_pro_:RR2b^Δ^*^54^–*GFP* and *35S_pro_:RR2b^Δ^*^137^–*GFP*) were transformed into *N. benthamiana* leaf epidermal cells for confocal microscopy. Scale bars = 20 μm. Three independent experiments were repeated with similar results. (f) Generation of *rr2b* knockout alleles in the c.v. W82 background by CRISPR– Cas9 gene editing. Two different edited events, *rr2b-1* (–1 bp) and *rr2b-2* (+1 bp), are shown for homozygous plants. (g) *RR2b* expression levels in transgenic over–expression lines OE 4 and OE 14 generated in the c.v. W82 background. Expression levels were normalized to *ELF1b* and data are presented as means ± SD from three biological replicates. Different letters indicate statistically significant differences at *p* < 0.05 by one-way ANOVA analysis with Tukey’s test.

**Figure S4.**
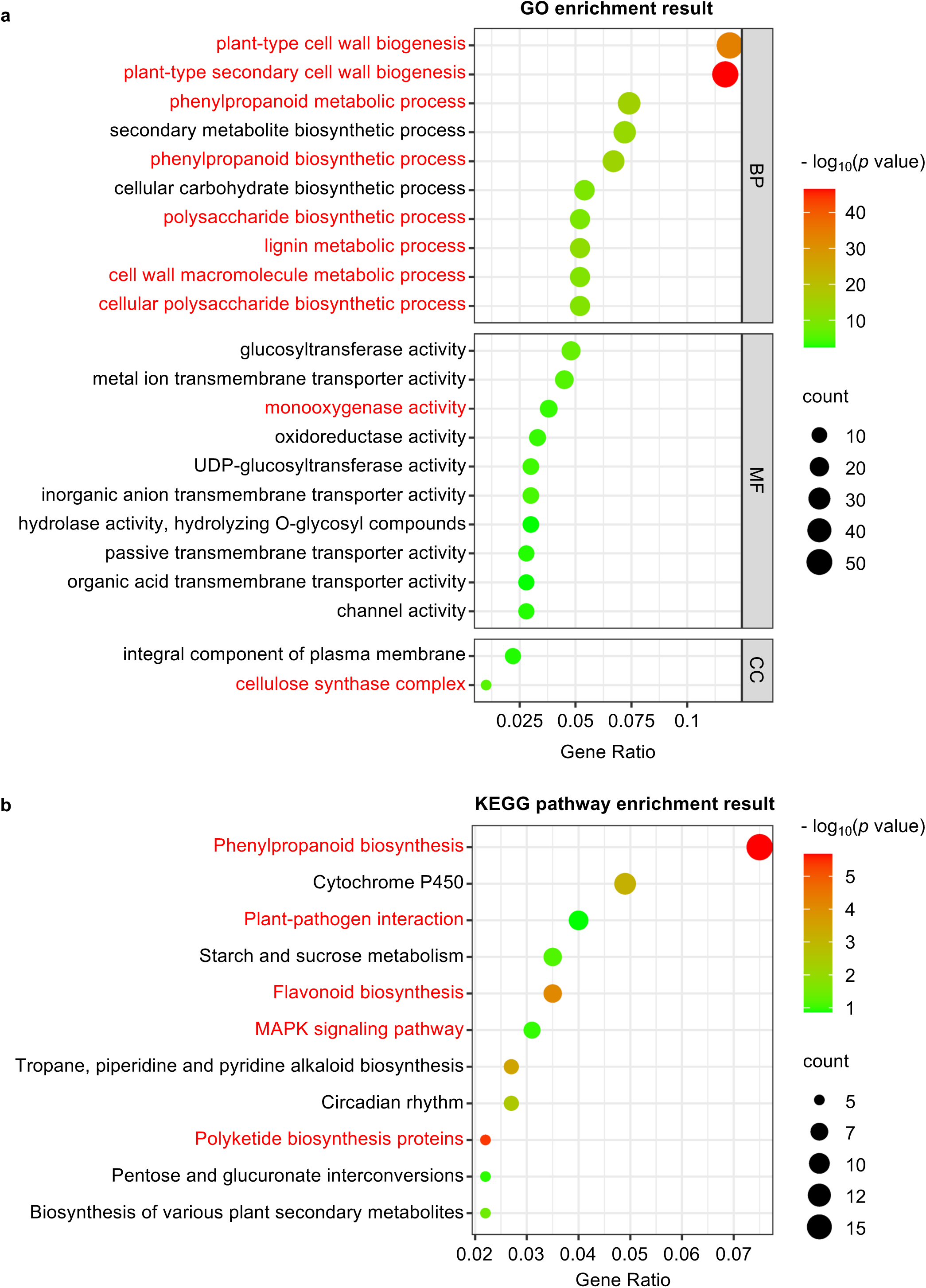
Cell-wall-biosynthesis and plant–pathogen-interaction-related pathways are enriched in *rr2b* knockout and *RR2b* over–expression backgrounds. (a) ClusterProfile was used to perform GO functional–enrichment analysis of the differential gene sets. Gene ontology (GO) is divided into three aspects: biological process (BP), molecular function (MF) and cellular component (CC). For GO functional enrichment, *p* < 0.05 was used as the threshold for statistically significant enrichment. (b) KEGG pathway enrichment analysis was performed as follows: Differentially expressed genes were selected based on the criteria of |log2(Fold Change)| ≥ 1 and an adjusted *p*-value (FDR) < 0.05. Enrichment significance of the target gene set in KEGG pathways was evaluated using a hypergeometric distribution test, with the entire genome– annotated gene set as the background reference. Benjamini–Hochberg testing was applied to correct for multiple hypothesis testing. Significantly enriched pathways were defined as those with an adjusted *p*-value < 0.05 and an enrichment factor > 1. Results were visualized via bubble plots to display the top-11 enriched pathways, annotated with corresponding gene counts. Red font denotes cell-wall-biosynthesis and plant–pathogen-interaction-related pathways.

**Figure S5.**
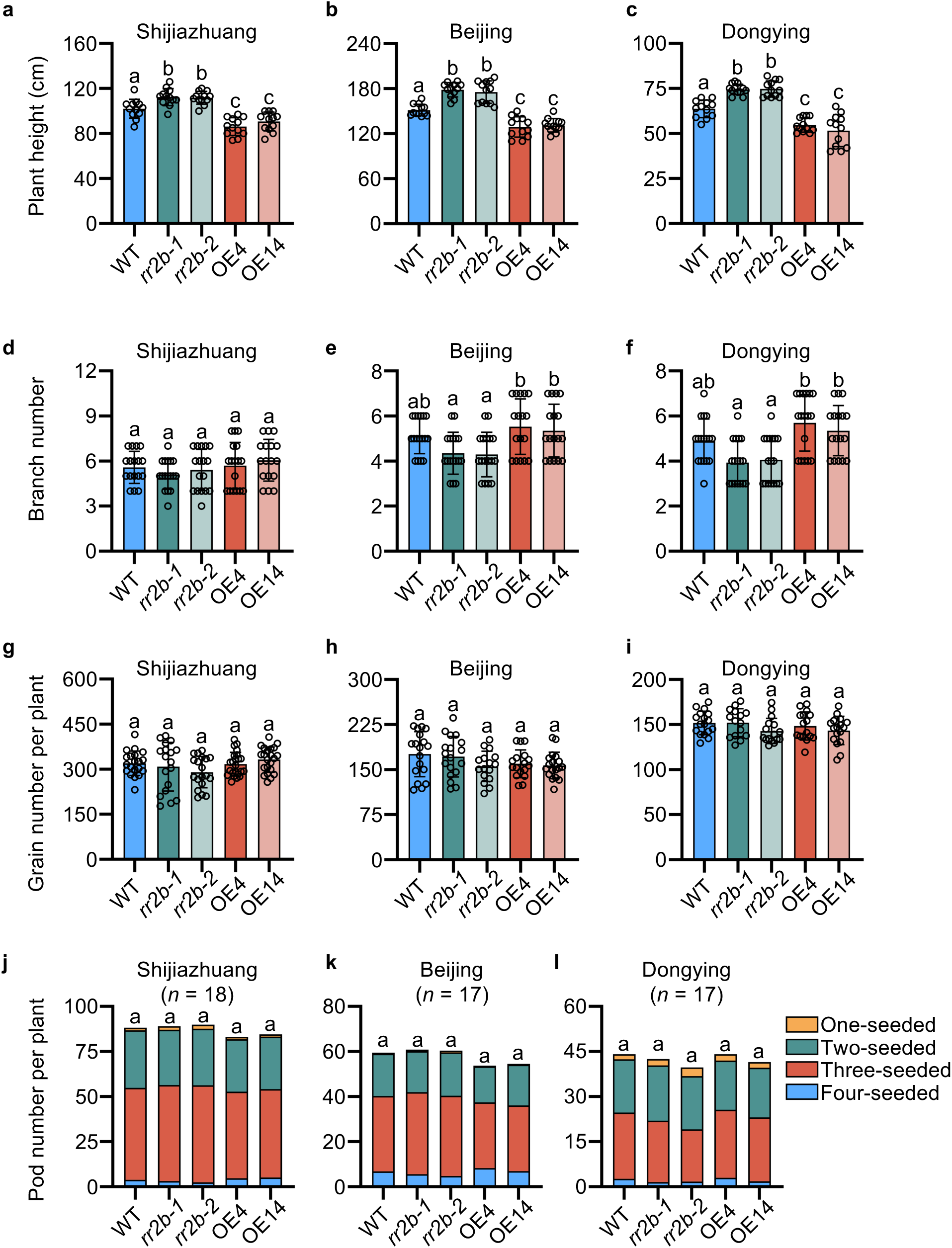
Over–expression and knockout of *RR2b* in soybean alter plant height but do not affect branch number, grain number per pod, and pod number in the field. (a–c) Plant height for W82, *rr2b* knockout and *RR2b* OE lines grown at Shijiazhuang (a), Beijing (b) and Dongying (c) field sites in 2024. (d–f) Branch numbers of W82, *rr2b* knockout and *RR2b* OE lines grown at multiple field sites in 2024. (g–i) Grain number per plant of W82, *rr2b* knockout and *RR2b* OE lines grown at multiple field sites in 2024. (j–l) Pod number per plant of W82, *rr2b* knockout and *RR2b* OE lines grown at multiple field sites in 2024. For a–i, data are presented as means ± SD (n = 18 individual plants) and different letters indicate statistically significant differences in a one-way ANOVA with Tukey’s *t*est (*p* < 0.05). For j–l, the number of biological replicates is indicated on the graph. Significance was analyzed by Fisher’s exact test compared with W82 (*p* < 0.05).

**Figure S6.**
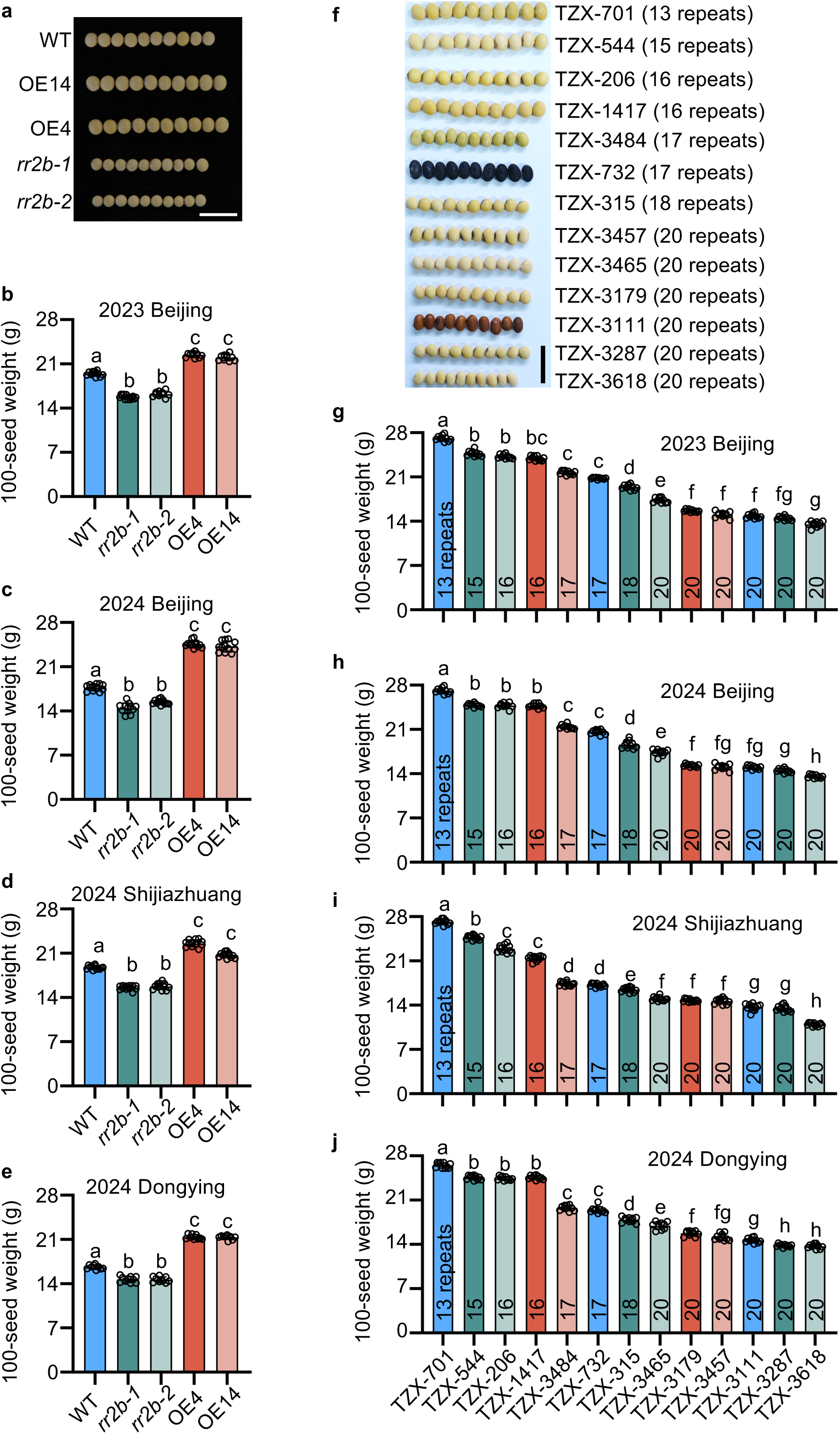
*RR2b* positively regulates soybean seed size. (a) Seeds size for W82, *rr2b* knockout and *RR2b* OE lines. Scale bars = 2 cm. (b–e) Quantification of 100-seed weight for W82, *rr2b* knockout and *RR2b* OE lines grown at multiple field sites in 2023 and 2024. (f) Seeds size of germplasm containing various insertion copies in the *RR2b* promoter. The numbers of insertion copies are listed in brackets. (g–j) Quantification of 100–seed weight for germplasm containing various insertion copies in the *RR2b* promoter grown at multiple field locations in 2023 and 2024. Data are presented as means ± SD and different letters indicate statistically significant differences at *p* < 0.05 by one-way ANOVA analysis with Tukey’s test. For g–j, the numbers on the bars refer to the copy numbers of (ATT) insertions.

**Figure S7.**
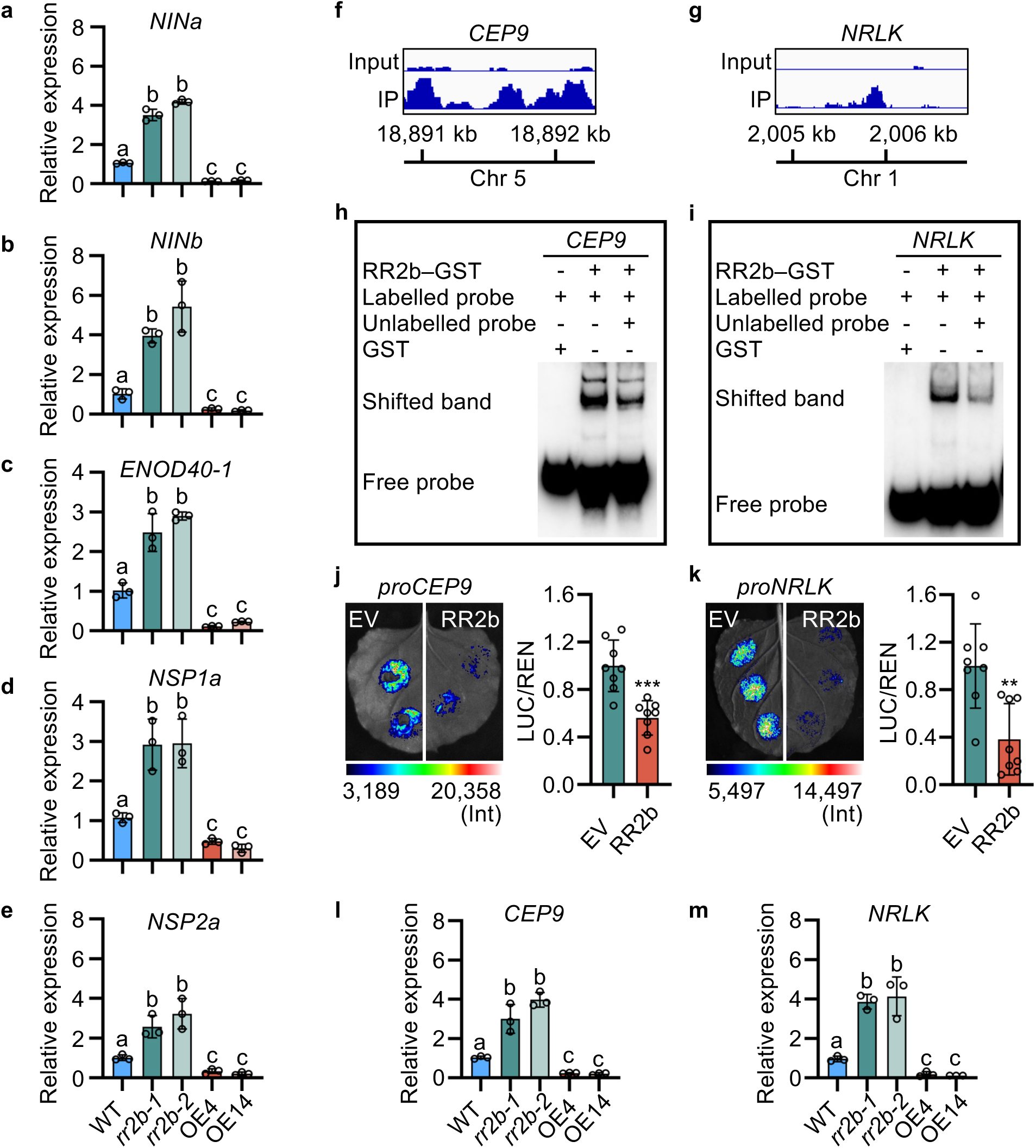
RR2b represses major soybean nodulation–pathway genes. (a–e) Relative expression of nodulation–pathway marker genes, *NINa* (a), *NINb* (b), *ENOD40-1* (c), *NSP1a* (d) and *NSP2a* (e) in W82, *rr2b* knockout and *RR2b* OE lines. Data are presented as means ± SD from three biological replicates. (f, g) CUT & Tag analysis of RR2b preferentially binding to the promoters of *CEP9* (f) and *NRLK* (g). (h, i) EMSA of GST–RR2b binding *in vitro* to *cis*– elements in the promoters of *CEP9* (h) and *NRLK* (i). Three independent replicates were performed and a representative result is shown. (j, k) Transient dual–luciferase assays of RR2b binding to the promoters of *CEP9* (j) and *NRLK* (k). Data are presented as means ± SD (n = 8). Three independent experiments were repeated with similar results. Asterisks indicate statistically significant differences relative to the EV control. Two–sided Student’s *t*–test, \*\**p* < 0.01, \*\*\**p* < 0.001. (l, m) Relative expression of *CEP9* (l) and *NRLK* (m) in W82, *rr2b* knockout and *RR2b* OE lines. Data are means ± SD from three biological replicates. In a–e, l and m, expression levels were normalized to *ELF1b*. Different letters indicate statistically significant differences at *p* < 0.05 by one-way ANOVA analysis with Tukey’s test.

**Figure S8.**
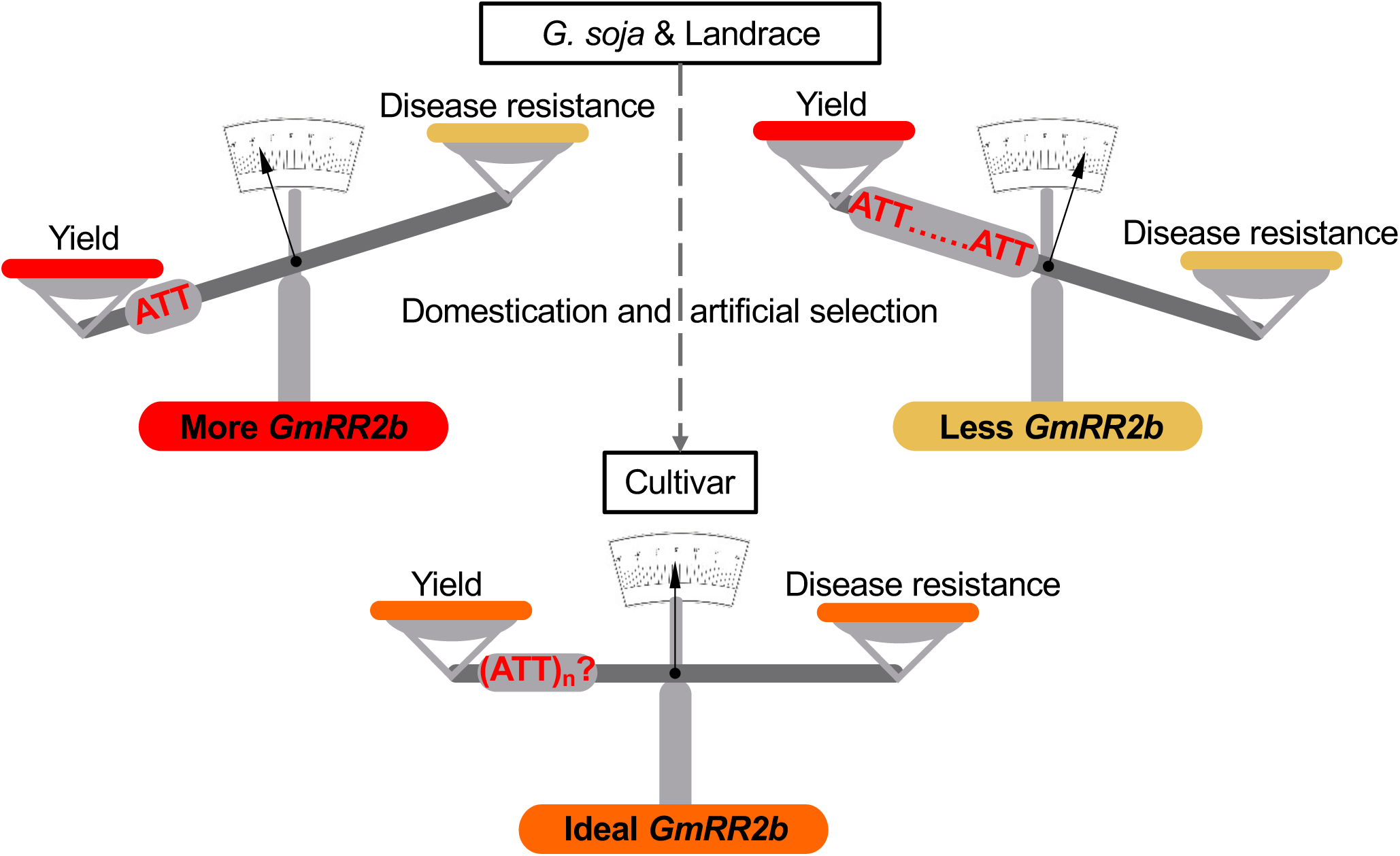
Proposed working model of RR2b function in balancing yield and disease resistance in soybean. In both wild soybeans and landraces, the *RR2b* promoter has varying lengths of ATT insertion fragments. Generally, longer insertion fragments correlate with weaker promoter activity and enhanced disease resistance in the corresponding varieties, but at the cost of reduced yield (the up-right balance). Conversely, shorter fragments tend to enhance yield but diminish disease resistance (the up-left balance). Throughout the soybean domestication process, haplotypes with moderate insertion–fragment lengths that facilitate optimal *RR2b* promoter activity were preserved, achieving a balance between yield and disease resistance (the bottom balance). The color of the balance pan indicates intensity levels: red signifies high intensity, yellow indicates low intensity, and orange denotes moderate intensity. The (ATT) insertion fragment represents the rider of the balance, precisely regulating soybean yield and disease resistance. The uncertain n and the question mark denote that the optimal *RR2b* activity, as determined by the length of the insertion fragment in its promoter, may vary under different environmental conditions for achieving the yield–disease resistance balance in soybeans

## References

1. Kim, M. Y., Van, K., Kang, Y. J., Kim, K. H. & Lee, S.-H. Tracing soybean domestication history: from nucleotide to genome. Breed. Sci. 61, 445–452 (2011).

2. FAO. FAOSTAT: crops and livestock products. (2023).

3. Ladha, J. K. et al. Biological nitrogen fixation and prospects for ecological intensification in cereal-based cropping systems. Field Crops Res. 283, 108541 (2022).

4. Hyten, D. L. et al. Impacts of genetic bottlenecks on soybean genome diversity. Proc. Natl. Acad. Sci. 103, 16666–16671 (2006).

5. Ross-Ibarra, J., Morrell, P. L. & Gaut, B. S. Plant domestication, a unique opportunity to identify the genetic basis of adaptation. Proc. Natl. Acad. Sci. 104, 8641–8648 (2007).

6. Frary, A. et al. *fw2.2*: a quantitative trait locus key to the evolution of tomato fruit size. Science 289, 85–88 (2000).

7. Li, C. B., Zhou, A. L. & Sang, T. Rice domestication by reducing shattering. Science 311, 1936–1939 (2006).

8. Doebley, J., Stec, A. & Gustus, C. Teosinte branched1 and the origin of maize: evidence for epistasis and the evolution of dominance. Genetics 141, 333–346 (1995).

9. Kato, K., Sonokawa, R., Miura, H. & Sawada, S. Dwarfing effect associated with the threshability gene *Q* on wheat chromosome 5A. Plant Breed. 122, 489–492 (2003).

10. Sun, L. et al. *GmHs1-1*, encoding a calcineurin-like protein, controls hard-seededness in soybean. Nat. Genet. 47, 939–943 (2015).

11. Wang, M. et al. Parallel selection on a dormancy gene during domestication of crops from multiple families. Nat. Genet. 50, 1435–1441 (2018).

12. Huang, X. et al. A map of rice genome variation reveals the origin of cultivated rice. Nature 490, 497–501 (2012).

13. Hufford, M. B. et al. Comparative population genomics of maize domestication and improvement. Nat. Genet. 44, 808–811 (2012).

14. Zhou, Z. et al. Resequencing 302 wild and cultivated accessions identifies genes related to domestication and improvement in soybean. Nat. Biotechnol. 33, 408–414 (2015).

15. Sakai, H., Aoyama, T. & Oka, A. *Arabidopsis* ARR1 and ARR2 response regulators operate as transcriptional activators. Plant J. 24, 703–711 (2000).

16. Zsigmond, L. et al. Overexpression of the mitochondrial *PPR40* gene improves salt tolerance in Arabidopsis. Plant Sci. 182, 87–93 (2012).

17. Zhang, Y., Ma, Y., Liu, R. & Li, G. Genome-wide characterization and expression analysis of KH family genes response to ABA and SA in *Arabidopsis thaliana*. Int. J. Mol. Sci. 23, 511 (2022).

18. Hao, Z. et al. Leucine-rich repeat protein family regulates stress tolerance and development in plants. Rice Science 32, 32–43 (2025).

19. Laffont, C. & Frugier, F. Rhizobium symbiotic efficiency meets CEP signaling peptides. New Phytol. 241, 24–27 (2024).

20. Laffont, C. et al. *MtNRLK1*, a CLAVATA1-like leucine-rich repeat receptor-like kinase upregulated during nodulation in *Medicago truncatula*. Sci. Rep. 8, 2046 (2018).

21. Liu, T., Longhurst, A. D., Talavera-Rauh, F., Hokin, S. A. & Barton, M. K. The *Arabidopsis* transcription factor ABIG1 relays ABA signaled growth inhibition and drought induced senescence. Elife 5, 13768 (2016).

22. Kurakawa, T. et al. Direct control of shoot meristem activity by a cytokinin-activating enzyme. Nature 445, 652–655 (2007).

23. Duan, J. et al. Strigolactone promotes cytokinin degradation through transcriptional activation of *CYTOKININ OXIDASE*/*DEHYDROGENASE 9* in rice. Proc. Natl. Acad. Sci. 116, 14319–14324 (2019).

24. Ashikari, M. et al. Cytokinin oxidase regulates rice grain production. Science 309, 741–745 (2005).

25. Wu, B. et al. Suppressing a phosphohydrolase of cytokinin nucleotide enhances grain yield in rice. Nat. Genet. 55, 1381–1389 (2023).

26. Jameson, P. E. & Song, J. Cytokinin: a key driver of seed yield. Journal of Experimental Botany 67, 593–606 (2016).

27. Argueso, C. T. & Kieber, J. J. Cytokinin: from autoclaved DNA to two-component signaling. Plant Cell 36, 1429–1450 (2024).

28. Hu, Y. et al. GmJAZ3 interacts with GmRR18a and GmMYC2a to regulate seed traits in soybean. J. Integr. Plant Biol. 65, 1983–2000 (2023).

29. Yang, Y. et al. Soybean type-B response regulator GmRR1 mediates phosphorus uptake and yield by modifying root architecture. Plant Physiol. 194, 1527–1544 (2024).

30. Alam, O. & Purugganan, M. D. Domestication and the evolution of crops: variable syndromes, complex genetic architectures, and ecological entanglements. Plant Cell 36, 1227–1241 (2024).

31. Zimmerer, K. S. & de Haan, S. Agrobiodiversity and a sustainable food future. Nat. Plants 3, 17047 (2017).

32. Ma, Y., Liu, M., Stiller, J. & Liu, C. A pan-transcriptome analysis shows that disease resistance genes have undergone more selection pressure during barley domestication. BMC Genomics 20, 12 (2019).

33. Burdon, J. J. & Thrall, P. H. The fitness costs to plants of resistance to pathogens. Genome Biol. 4, 227 (2003).

34. Barabaschi, D., Tondelli, A., Vale, G. & Cattivelli, L. Fitness cost shapes differential evolutionary dynamics of disease resistance genes in cultivated and wild plants. Mol. Plant 13, 1352–1354 (2020).

35. Cordova-Campos, O., Adame-Alvarez, R. M., Acosta-Gallegos, J. A. & Heil, M. Domestication affected the basal and induced disease resistance in common bean (*Phaseolus vulgaris*). Eur. J. Plant Pathol. 134, 367–379 (2012).

36. Varshney, R. K. et al. Resequencing of 429 chickpea accessions from 45 countries provides insights into genome diversity, domestication and agronomic traits. Nat. Genet. 51, 857–864 (2019).

37. Mittler, R. ROS are good. Trends Plant Sci. 22, 11–19 (2017).

38. Choi, J. et al. The cytokinin-activated transcription factor ARR2 promotes plant immunity via TGA3/NPR1-dependent salicylic acid signaling in *Arabidopsis*. Dev. Cell 19, 284–295 (2010).

39. Arnaud, D. et al. Cytokinin-mediated regulation of reactive oxygen species homeostasis modulates stomatal immunity in Arabidopsis. Plant Cell 29, 543–559 (2017).

40. Tan, S. et al. A cytokinin signaling type-B response regulator transcription factor acting in early nodulation. Plant Physiol. 183, 1319–1330 (2020).

41. Chen, J. et al. The B-type response regulator GmRR11d mediates systemic inhibition of symbiotic nodulation. Nat. Commun. 13, 7661 (2022).

42. Hung, H. Y., et al. *ZmCCT* and the genetic basis of day-length adaptation underlying the postdomestication spread of maize. Proc. Natl. Acad. Sci. 109, E1913–E1921 (2012).

43. Liu, C. et al. Early selection of *bZIP73* facilitated adaptation of *japonica* rice to cold climates. Nat. Commun. 9, 3302 (2018).

44. Li, C. et al. A domestication-associated gene *GmPRR3b* regulates the circadian clock and flowering time in soybean. Mol. Plant 13, 745–759 (2020).

45. Lu, S. et al. Stepwise selection on homeologous *PRR* genes controlling flowering and maturity during soybean domestication. Nat. Genet. 52, 428–436 (2020).

46. He, Z., Webster, S. & He, S. Y. Growth-defense trade-offs in plants. Curr. Biol. 32, R634–R639 (2022).

47. Wu, W. et al. A single-nucleotide polymorphism causes smaller grain size and loss of seed shattering during African rice domestication. Nat. Plants 3, 17064 (2017).

48. Li, S. et al. Modulating plant growth-metabolism coordination for sustainable agriculture. Nature 560, 595–600 (2018).

49. Wang, J. et al. A single transcription factor promotes both yield and immunity in rice. Science 361, 1026–1028 (2018).

50. Song, X. et al. Targeting a gene regulatory element enhances rice grain yield by decoupling panicle number and size. Nat. Biotechnol. 40, 1403–1411 (2022).

51. Jian, B. et al. Validation of internal control for gene expression study in soybean by quantitative real-time PCR. BMC Mol. Biol. 9, 59 (2008).

52. Danecek, P. et al. The variant call format and VCFtools. Bioinformatics 27, 2156–2158 (2011).

53. Liu, Y., et al. Pan-Genome of Wild and Cultivated Soybeans. Cell 182, 162–176 (2020).

54. Liang, Q. et al. Natural variation of *Dt2* determines branching in soybean. Nat. Commun. 13, 6429 (2022).

55. Sun, B. et al. A high-resolution transcriptomic atlas depicting nitrogen fixation and nodule development in soybean. J. Integr. Plant Biol. 65, 1536–1552 (2023).

56. Ma, X. et al. A robust CRISPR/Cas9 system for convenient, high-efficiency multiplex genome editing in monocot and dicot plants. Mol. Plant 8, 1274–1284 (2015).

57. Lei, Y. et al. CRISPR-P: a web tool for synthetic single-guide RNA design of CRISPR-system in plants. Mol. Plant 7, 1494–1496 (2014).

58. Ji, H. et al. Differential light-dependent regulation of soybean nodulation by papilionoid-specific HY5 homologs. Curr. Biol. 32, 783–795 (2022).

59. Ren, Z. et al. The BRUTUS iron sensor and E3 ligase facilitates soybean root nodulation by monoubiquitination of NSP1. Nat. Plants 11(2025).

60. Burnette, W. N. “Western blotting”: electrophoretic transfer of proteins from sodium dodecyl sulfate-polyacrylamide gels to unmodified nitrocellulose and radiographic detection with antibody and radioiodinated protein A. Analytical Biochemistry 112, 195–203 (1981).

61. Yu, H. et al. GmNAC039 and GmNAC018 activate the expression of cysteine protease genes to promote soybean nodule senescence. Plant Cell 35, 2929–2951 (2023).

62. Yoo, C. Y. et al. The *Arabidopsis* GTL1 transcription factor regulates water use efficiency and drought tolerance by modulating stomatal density via transrepression of *SDD1*. Plant Cell 22, 4128–4141 (2010).

63. Li, L. et al. The FLS2-associated kinase BIK1 directly phosphorylates the NADPH oxidase RbohD to control plant immunity. Cell Host Microbe 15, 329–338 (2014).

64. Wu, X. et al. SNP discovery by high-throughput sequencing in soybean. BMC Genomics 11, 469 (2010).

65. Abdelmajid, K. M. et al. Quantitative trait loci (QTL) that underlie SCN resistance in soybean [*Glycine max* (L.) Merr.] PI438489B by ‘Hamilton’ re-combinant inbred line (RIL) population. Atlas J. Plant Biol. 1, 29–38 (2014).

66. Ning, H. et al. Identification of QTLs related to the vertical distribution and seed-set of pod number in soybean *Glycine max* (L.) Merri. PLoS One 13, e0195830. (2018).

67. Luckew, A. S., Swaminathan, S., Leandro, L. F., Orf, J. H. & Cianzio, S. R. ’MN1606SP’ by ’Spencer’ filial soybean population reveals novel quantitative trait loci and interactions among loci conditioning SDS resistance. Theor. Appl. Genet. 130, 2139–2149 (2017).

68. Harris, D. K. et al. Soybean quantitative trait loci conditioning soybean rust-induced canopy damage. Crop Sci. 55, 2589–2597 (2015).

69. Zhang, D. et al. Identification of genomic regions determining flower and pod numbers development in soybean (*Glycine max* L.). J. Genet. Genomics 37, 545–556 (2010).

70. Wang, Y. et al. Mapping isoflavone QTL with main, epistatic and QTL x environment effects in recombinant inbred lines of soybean. PLoS One 10, e0118447 (2015).

71. Han, Y. et al. Unconditional and conditional QTL underlying the genetic interrelationships between soybean seed isoflavone, and protein or oil contents. Plant Breed. 134, 300–309 (2015).

